# Enhanced generalization and specialization of brain representations of semantic knowledge in healthy aging

**DOI:** 10.1101/2023.08.08.552519

**Authors:** Pedro Margolles, David Soto

**Author notes:** **Author Note:** Correspondence should be addressed to, Basque Center on Cognition, Brain and Language, Paseo Mikeletegi 69, 2nd Floor 20009 San Sebastian.

## Abstract

Aging is often associated with a decrease in cognitive capacities. However, semantic memory appears relatively well preserved in healthy aging. Both behavioral and neuroimaging studies support the view that changes in brain networks contribute to this preservation of semantic cognition. However, little is known about the role of healthy aging in the brain representation of semantic categories. Here we used pattern classification analyses and computational models to examine the neural representations of living and non-living word concepts. The results demonstrate that brain representations of animacy in healthy aging exhibit increased similarity across categories, even across different task contexts. This pattern of results aligns with the neural dedifferentiation hypothesis that proposes that aging is associated with decreased specificity in brain activity patterns and less efficient neural resource allocation. However, the loss in neural specificity for different categories was accompanied by increased dissimilarity of item-based conceptual representations within each category. Taken together, the age-related patterns of increased generalization and specialization in the brain representations of semantic knowledge may reflect a compensatory mechanism that enables a more efficient coding scheme characterized by both compression and sparsity, thereby helping to optimize the limited neural resources and maintain semantic processing in the healthy aging brain.

## Introduction

Aging is typically associated with a decline in performance across different cognitive domains including executive functions (Turner & Spreng, 2012), memory (Balota et al., 2000), attention (Madden, 2007) and general speed of processing (Salthouse, 2000), thereby impacting our quality of life. However, contrary to the decline observed in various cognitive domains, research has consistently shown that semantic memory tends to be relatively well preserved in healthy aging.

The preservation of semantic memory in aging can be attributed to the accumulation of knowledge and experiences acquired through lifetime. As individuals age, their conceptual richness and vocabulary size tend to remain stable or even improve (Rönnlund et al., 2005; Spaniol et al., 2006; Verhaeghen, 2003). Notably, behavioral studies observed that an expanded vocabulary is also associated with global modifications in the structure of semantic memory networks (Kenett & Thompson-Schill, 2020; Wulff et al., 2019; Yee & Thompson-Schill, 2016). These alterations may render the semantic networks less flexible and efficient, becoming more segregated and modular compared to younger individuals (Cosgrove et al., 2021, 2023; Dubossarsky et al., 2017; Wulff et al., 2019, 2022), thereby leading to changes in semantic knowledge use (e.g., Brosseau & Cohen, 1996; Kintz & Wright, 2017; Morrow & Duffy, 2005; Verheyen et al., 2019). Moreover, previous studies have also indicated that compensatory brain mechanisms contribute to maintaining semantic cognition in aging. Binder et al. (2009) identified increased left hemisphere activity during semantic tasks, a pattern observed in healthy young adults. However, Hoffman and Morcom (2018) discovered that older adults shift from a left-lateralized semantic network to a more evenly distributed bilateral activation pattern. Additionally, they observed a transition from domain-specific to domain-general neural resources in this older demographic.

Despite significant progress in understanding the relatively preserved nature of semantic memory in aging and the compensatory brain mechanisms in language and domain-general cognition, there remains a significant gap in our understanding of how healthy aging shapes the brain representation of semantic knowledge. The previous neuroimaging studies have primarily relied on univariate measures of brain activation. However, these measures, which average activity across individual voxels within a specific brain region, overlook the voxel covariance and the multivariate nature of brain activity patterns, potentially resulting in cases where distinct mental states generate indistinguishable univariate activation (Hebart & Baker, 2018; Ritchie et al., 2019). In contrast, multivariate pattern analysis (MVPA) methods (Haxby et al., 2001; Haynes & Rees, 2006; Norman et al., 2006; Tong & Pratte, 2012) offer higher specificity and sensitivity when exploring information encoded within distributed patterns of neural activity. Recent MVPA studies have established a link between aging and a decline in the distinctiveness of brain activity patterns, coupled with a diminished ability to efficiently allocate neural resources according to task goals (neural dedifferentiation hypothesis). Several factors likely contribute to age-related neural dedifferentiation, including variations in neuromodulatory drive, the effectiveness of inhibitory neurotransmission, responsiveness to task demands, and the cumulative impact of life experiences (Koen & Rugg, 2019). Two significant MVPA techniques for exploring neural changes associated with aging include standard machine learning linear classifiers to categorize different stimuli (e.g., Park et al., 2010; Carp, Park, Polk, et al., 2011), and conducting model-based representational similarity analysis (RSA) (e.g., Zheng et al., 2018). RSA is particularly valuable as it evaluates the similarity of neural activity patterns between stimuli within the same category and those across different categories (Zheng et al., 2018). This method not only enables researchers to discern how neural representations of stimuli evolve across different conditions, but also facilitates comparability with computational models acting as benchmarks, thereby shedding light on the underlying neural structure implicated in cognitive processes.

In this study, we utilize both machine learning pattern classifiers and RSA analyses, drawing from linguistic and computer vision models. Our aim is to examine the effects of aging on the brain’s representation of semantic categories associated with animacy, focusing on words related to living and non-living objects. Additionally, this work addresses the role of the depth of processing in the brain representation of semantic knowledge. Previous neuroimaging studies in healthy young adults have primarily focused on examining how the brain represents semantic knowledge by using tasks in which participants had either to attend to the items (Buchweitz et al., 2012; Shinkareva et al., 2011; Simanova et al., 2014), think about the stimuli (Mitchell et al., 2008), or make semantic judgments (Fernandino et al., 2016). However, the influence of task demands on the neural representation of semantic knowledge remains to be fully elucidated (Soto et al., 2020). In this study, the depth of processing was operationalized employing two conditions. The first condition involved silent reading, which required covertly reading the words and represented shallow processing. The second condition involved mental simulation of the concepts’ properties, which required actively imagining sensory-specific representations, representing a deeper level of processing. This condition required a more profound comprehension of word meanings compared to the shallow processing, which primarily focused on phonological aspects. For further details on the experimental procedure, refer to the Methods section and Figure 1. Grasping how increased task demands affect the brain’s representation of semantic categories is vital for understanding the neurocognitive impacts of aging. In response to these demands, older adults may employ compensatory mechanisms to maintain cognitive performance (Hoffman & Morcom, 2018). These mechanisms are particularly important under deep processing conditions that require more cognitive resources (Haitas et al., 2021; Reuter-Lorenz & Cappell, 2008). Age-related declines in physical and sensorimotor abilities can further challenge older adults by disrupting their ability to integrate information needed for concept formation and mental simulations (Costello & Bloesch, 2017; Kuehn et al., 2018; Mille et al., 2021; Vallet, 2015). These conditions may have implications for the brain’s representation of semantic knowledge in older adults relative to younger counterparts.

**Figure 1.**
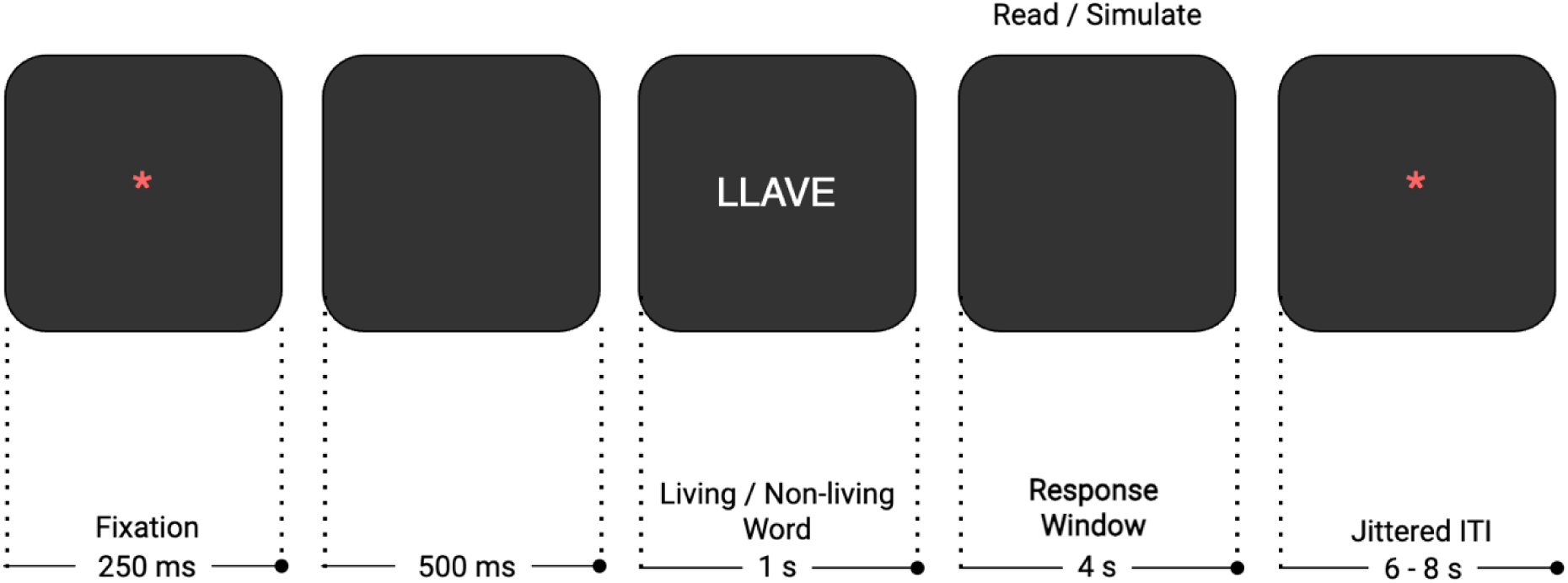
Illustration of an experimental trial in both depth of processing conditions (i.e., shallow, and deep).

## Experimental Overview

Our initial goal was to explore how the distinctiveness of brain activity patterns associated with semantic categories changes with aging in the bilateral semantic network. We conducted MVPA decoding, utilizing a machine learning classifier to determine the semantic categories of previously unseen living and non-living words, in both within and across conditions. The decoding accuracy of our classifier is the performance metric we used to evaluate the distinctiveness of these semantic categories. In within-condition analyses, we anticipated that decoding accuracy would be lower in older adults compared to younger adults, attributed to the neural dedifferentiation of semantic representations. This effect is expected to be more pronounced in the deep processing condition, which demands greater cognitive resources and sensorimotor abilities. Moreover, these demands could affect the integration of semantic representations, potentially leading to compensatory patterns where decoding accuracy is reduced in the left hemisphere and increased in the right hemisphere. In cross-condition analyses, we hypothesized that if aging affects the generalization of brain representations across different tasks, making them less specific but more generalizable, then we would expect superior classification performance in this group compared to young adults. Next, we investigated the neural representations of living and non-living word categories across different age groups. Using searchlight RSA, we compared these neural structures with those derived from a language model based on word embeddings, and a computer vision model fed with image referents associated with the concepts. This RSA approach enabled us to gain a comprehensive understanding of the similarities and differences in brain representation of animacy categories across age groups.

## Methods

### Participants

Twenty-seven young adults from a previous study (Soto et al., 2020), and 29 healthy older adults participated in the research and received monetary compensation in return. The experimental procedures were carried out following the principles stated in the Declaration of Helsinki and received approval from the BCBL Research Ethics Board. Written informed consent was obtained from all participants. Refer to Table 1 below for sociodemographic and neuropsychological characteristics of the sample.

**Table 1.** Sociodemographic and neuropsychological characteristics of experimental groups. To ensure that the level of cognitive performance of older adults fell within the normal range for their age, a neuropsychological assessment was administered using standardized instruments adapted for the Spanish population: Mini-Mental State Examination (MMSE; Folstein et al., 1975; Lobo et al., 1980), Montreal Cognitive Assessment (MoCA; Nasreddine et al., 2005), Repeatable Battery for the Assessment of Neuropsychological Status (RBANS; Randolph et al., 1998). Additionally, the cognitive reserve of older adults, which assesses their engagement in cognitively stimulating activities over their lifetimes, was measured using the Cognitive Reserve Questionnaire (CRQ; Rami et al., 2011) and the Cognitive Reserve Scale (CRS; León et al., 2011, 2014, 2016).

|  |  | Mean (Reference values) | SD |
| --- | --- | --- | --- |
| Age | Young group (n=27, 10 males) | 24 | 3 |
|  | Old group (≥65 years old; n=29, 12 males) | 71 | 4 |
| Language | Native spanish speakers |  |  |
| Handedness | All right-handed |  |  |
| Vision | Normal or corrected-to-normal |  |  |
| Examinations | Normal physical and neurological examinations |  |  |
| Neuropsychological Assessment of Older Adults |  |  |  |
| Cognitive Impairment | MMSE | 29.21 (≥27 normal) | 1.01 |
|  | MoCA | 26.66 (≥26 normal) | 1.91 |
|  | RBANS (Global Index) | Global Index: 112.31 (percentile: 76) | Global Index: 13.98 Percentile: 24.29 |
| Cognitive Reserve | CRQ | 16.9 (≥15 high reserve) | 3.75 |
|  | CRS (Global Score) | 173,66 (percentile: 79) | 31,70 |
| Sociodemographic Assessment of Older Adults |  |  |  |
| Educational attainment |  | Basic studies (6 participants)<br>Mid studies (12 participants)<br>High studies (11 participants) |  |
| Occupational attainment |  | Low (9 participants)<br>Mid (12 participants)<br>High (8 participants) |  |

### MRI settings

Functional images were acquired from each participant using a SIEMENS Magnetom Prisma-fit whole-body MRI scanner with a 3T magnet and a 64-channel head coil. During each fMRI run, a multiband gradient-echo echo-planar imaging sequence was employed to capture 530 3D volumes encompassing the entire brain. The sequence incorporated a multi-band acceleration factor of 6, resulting in a voxel resolution of 2.4 * 2.4 * 2.4 mm³. The acquisition parameters included a time-to-repetition of 850 ms, a time-to-echo of 35 ms, and a bandwidth of 2582 Hz/Px. The acquired volumes covered 66 slices with a field of view of 210. After the completion of the initial four functional runs, a high-resolution T1-weighted structural MPRAGE scan was obtained with a voxel size of 1 * 1 * 1 mm³.

### Experimental task

The experimental procedure consisted of 8 fMRI runs (≈ 8 minutes each), with a 1-2-minute break between each run. At the beginning of each trial, a red fixation asterisk was displayed for 250 ms, followed by a 500 ms blank screen. In each run, participants completed 36 trials, with each trial involving a randomly chosen Spanish word from a set of 18 living and 18 non-living words. Table S1 in the supplementary materials provides a breakdown of the experimental words by category, along with the characteristics of living and non-living words.

The selected word appeared at the screen’s center for 1 second, followed by a 4-second response window (see Figure 1). Participants were instructed to either covertly read and mentally repeat the word (shallow processing condition) or engage in mental simulations considering the sensorimotor properties like shape, color, sound, and context of the represented concept (deep processing condition). The two conditions alternated between runs. In order to improve the signal-to-noise ratio (SNR) in MVPA analyses, each word was shown 4 times per condition (once per run). In the deep processing condition, participants were instructed to employ similar mental simulations when encountering the same word again.

To ensure that the BOLD response was separated between trials, the interstimulus interval period was jittered between 6 and 8 seconds and indicated by a red asterisk. The duration of the ITI was determined using a pseudo-exponential distribution, resulting in different intervals for each trial. Specifically, 50% of the trials had a 6-second interval, 25% had a 6.5-second interval, 12.5% had a 7-second interval, and so on. To maintain participants’ attention, up to two catch trials were randomly inserted in each run, consistent across both tasks. These trials replaced living and non-living words with number words (e.g., CERO, UNO, TRES), prompting participants to press any of the four buttons on an fMRI-compatible response pad.

### MVPA analysis

In this section, we present a detailed overview of our MVPA approach, which provides enhanced specificity and sensitivity in probing the information encoded within distributed patterns of neural activity. Our MVPA analyses aim to investigate the neural changes associated with aging, particularly focusing on alterations in the brain’s representation of semantic categories linked to animacy. Additionally, we explore the influence of depth of processing on semantic representations using decoding analyses with a machine learning pattern classifier, and searchlight RSA based on computational models. Through this integrative approach, we aim to unravel the intricate neural dynamics underlying age-related changes in semantic cognition, shedding light on how aging influences the organization of semantic knowledge in the brain. A figure illustrating a comprehensive overview of the MVPA pipeline can be found in Supplementary materials (Figure S1).

### MRI data preprocessing

A sequence of preprocessing steps was implemented on each fMRI run utilizing FSL (Jenkinson et al., 2012; S. M. Smith et al., 2004) and its FEAT tool (Woolrich et al., 2001, 2004). To ensure image stabilization, the first 9 volumes were discarded, and head-motion was corrected using MCFLIRT (Jenkinson et al., 2002). Non-brain voxels were removed via functional brain extraction, followed by minimal spatial smoothing using a 3 mm FWHM Gaussian Kernel. Runs 2-8 were co-aligned to a reference volume from the first preprocessed run. Motion-induced variations were corrected using ICA-AROMA (Pruim et al., 2015), and high-pass temporal filtering was applied with a 60-second cutoff to refine the functional data using FEAT.

### Decoding analyses

Our initial goal was to investigate the changes in the distinctiveness of brain activity patterns associated with semantic categories as people age, focusing on the bilateral semantic network. We utilized a machine learning classifier to predict the semantic categories of unseen experimental words. Our analysis, applied within and across two conditions that varied in processing depth, sought to assess these changes in distinctiveness with aging.

Drawing from a prior meta-analysis of the semantic system (Binder et al., 2009), we extracted a predefined set of 15 regions of interest (ROIs) for each hemisphere (see Figure 2). The extraction process utilized the FreeSurfer v.6.0 recon-all brain reconstruction algorithm, which was applied to the structural image (Fischl, 2012). The mri_binarize command from FreeSurfer was used to create anatomical masks for 15 ROIs by extracting cortical label indices from the aparc + aseg file and matching them with entries in the FreeSurferColorLUT text file (surfer.nmr.mgh.harvard.edu/fswiki/FsTutorial/AnatomicalROI/FreeSurferColorLUT). These masks were adjusted to each preprocessed run’s functional space using FLIRT in FSL with a 7 degrees-of-freedom (DOF) linear registration, and finally binarized.

**Figure 2.**
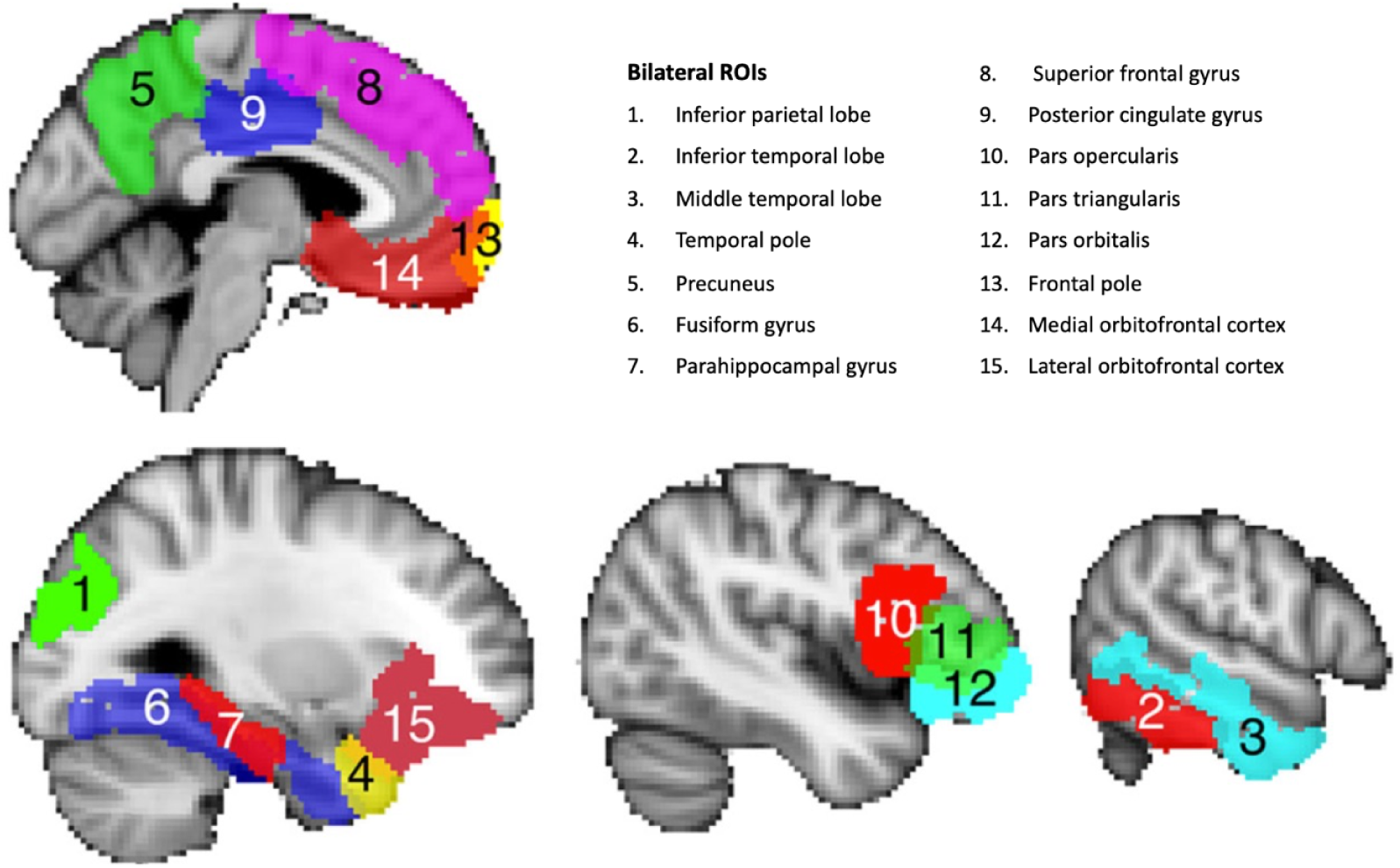
Illustration of the anatomical ROIs used in the decoding analyses based on Binder et al. (2009) meta-analysis.

MVPA decoding was conducted using Python 2.7, utilizing Scikit-learn (Pedregosa et al., 2011) and PyMVPA (Hanke et al., 2009). Functional data within each ROI were extracted using binarized masks, applied across all fMRI runs with PyMVPA. Psychopy-generated behavioral data provided labels for time points within each preprocessed fMRI run mask (word, category, condition…).

After stacking data from all functional runs separately for each ROI, invariant features were removed to prevent numerical issues during Z-scoring. Data underwent run-wise linear detrending and Z-score standardization. To account for the hemodynamic lag and enhance the SNR for decoding, a single instance was created for each individual trial by averaging voxel intensities from 3.4s to 8.6s after the onset of the word (≈ 6 volumes). To assess whether brain activation patterns contained semantic category information (living vs. non-living), we utilized a linear support vector machine (SVM) classifier with default parameters from Scikit-learn (l2 regularization, C = 1.0, tolerance = 0.0001) in both within-condition and cross-condition generalization scenarios.

For within-condition analysis, data were divided into balanced training and test sets, with each training set containing instances of 34 concepts and the test set including one living and one non-living concept instances. In cross-condition analysis, a similar method was employed (Kaplan et al., 2015), where a SVM was trained on 34 concepts from one condition and tested on the remaining 2 concepts from another condition (i.e., from shallow to deep and vice versa). To reduce overfitting and dimensionality, Principal Component Analysis (PCA) was applied to the training set, and then projected onto the test set. The number of components from PCA was tailored to match the instances in the training set, ensuring equal representation across ROIs (Mourão-Miranda et al., 2005). The resulting PCA features were used to train the decoding model. Decoding performance was evaluated on the model accuracy in predicting the semantic categories of unseen instances, both within the same condition and across different conditions. Leave-pair-words-out cross-validation method was applied covering all combinations of training and test sets. Decoding accuracies were averaged across 324 folds for every condition/cross-condition direction, participant, and ROI. Refer to Figure 3 for within-condition decoding and Figure 4 for cross-condition decoding results.

**Figure 3.**
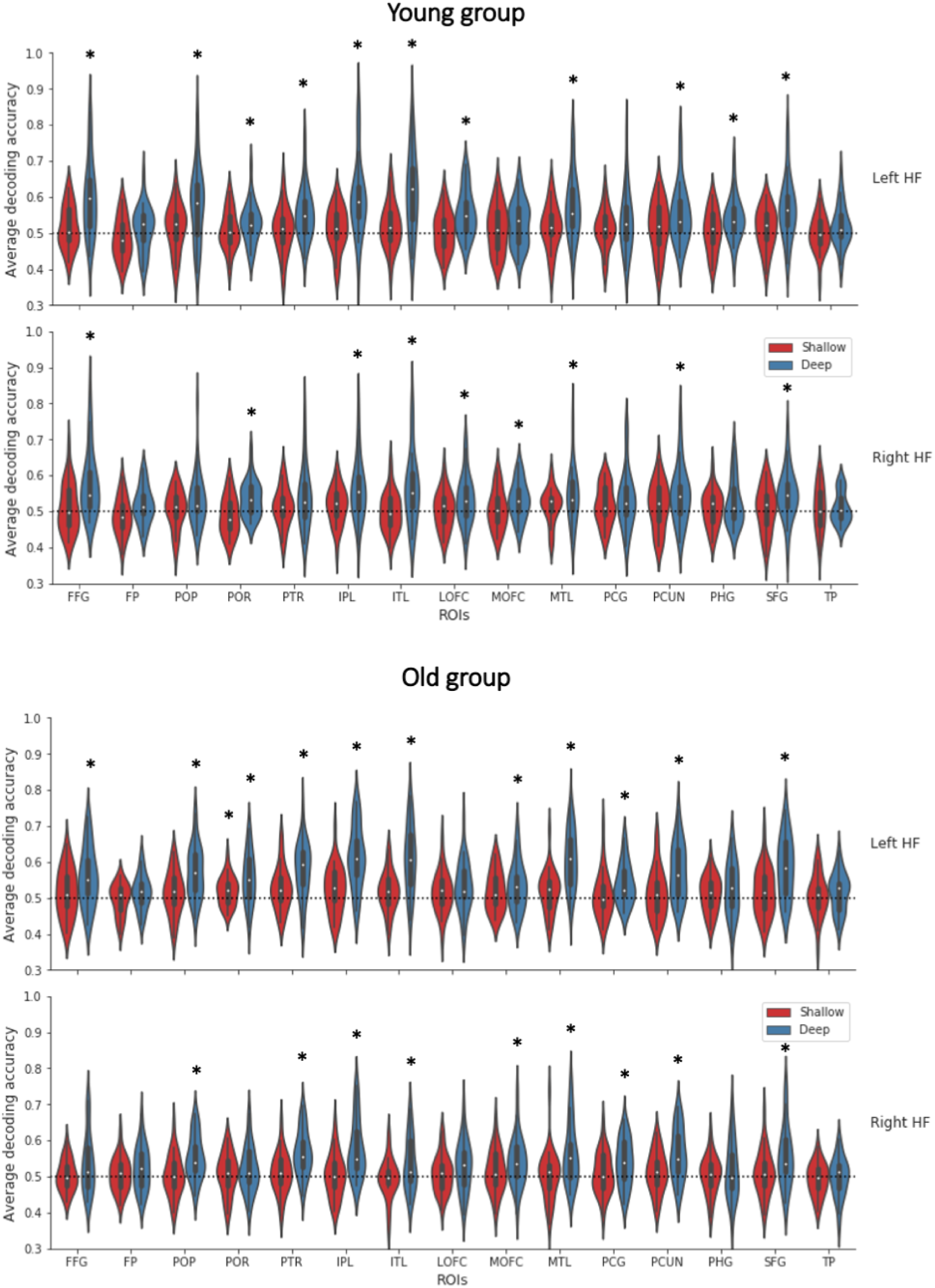
Within-condition decoding accuracies by ROI, condition, hemisphere, and age-group represented with violin plots. The black dotted line represents chance level (0.5). ROIs in which decoding performance was significantly above chance within each group in a one-sample t-test (FDR corrected p-values) are marked with an asterisk.

**Figure 4.**
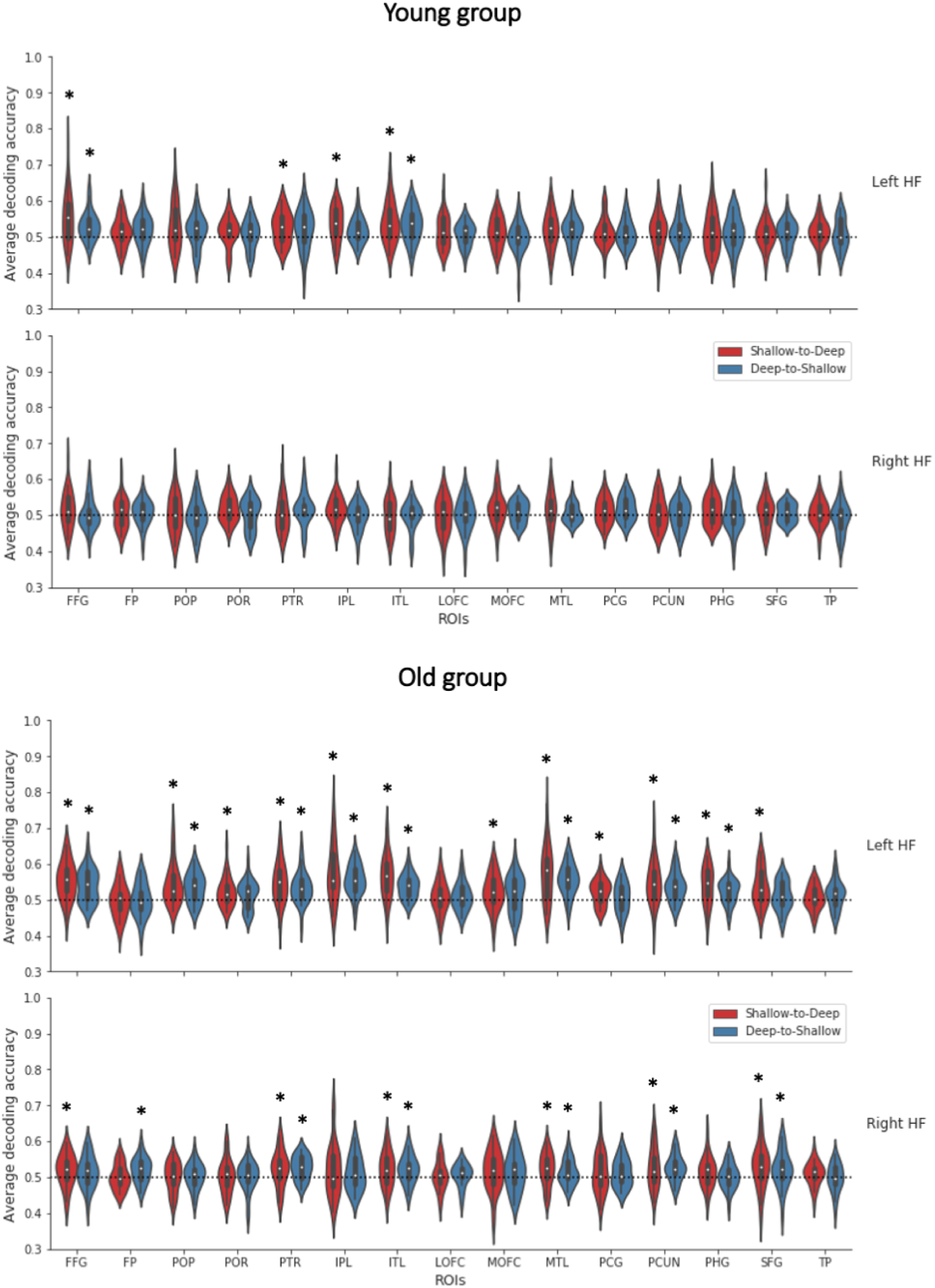
Cross-condition decoding accuracies by ROI, cross-condition direction, hemisphere, and age-group represented with violin plots. The black dotted line represents chance level (0.5). ROIs in which decoding performance was significantly above chance within each group in a one-sample t-test (FDR corrected p-values) are marked with an asterisk.

Finally, we conducted two mixed effects ANOVAs to compare the decoding accuracy across various factors in both within and cross-condition contexts. The ANOVAs encompassed 15 ROIs, 2 conditions (shallow vs. deep, or shallow-to-deep vs. deep-to-shallow), 2 hemispheres (left vs. right), and 2 age groups (older vs. young). Subsequent post-hoc pairwise comparisons were conducted utilizing the Python Scipy and Statsmodels modules (Seabold & Perktold, 2010; Virtanen et al., 2020) to explore main and interaction effects highlighted in the ANOVAs. P-values in the pairwise comparisons underwent correction for multiple comparisons employing the False Discovery Rate (FDR) method.

### Searchlight RSA

Next, we conducted an RSA analysis that compared brain patterns of both age groups with representations of experimental words derived from two computational models. By quantifying the similarity between distributed neural patterns and predefined computational models, searchlight RSA can illuminate how specific semantic categories are encoded across different regions of the brain as individuals age, highlighting the relationship between human cognitive processing and algorithmic predictions.

To investigate age-related differences in how semantic categories are organized in the brain, we utilized two advanced computational models. First, a Word2Vec model was used to assess semantic similarities among word categories (Cardellino, 2016). This model helps in understanding the linguistic relationships and clustering of words within different semantic fields. Second, we employed the VGG-19 model, a deep learning tool for image processing (Simonyan & Zisserman, 2014), to compare features from images related to the living/non-living concepts (e.g., Anderson et al., 2015, 2019; Deng et al., 2021; Devereux et al., 2018; Martin et al., 2018). The inclusion of a computer vision model was motivated by the cognitive demands of the mental simulation condition, which required participants to vividly simulate the sensory properties of each concept. We synthesized this data into a standardized representational dissimilarity matrix per model, using Pearson correlation distance to quantify the dissimilarity among word pairs based on computational features. A detailed explanation of the feature extraction process for each model is available in the Supplementary materials. This approach enabled us to systematically analyze and compare models’ representations with brain representations across different ages using RSA.

In this context, we processed the whole-brain raw data using the same pipeline employed for ROI-based decoding. Searchlight RSA analyses for each experimental condition and computer model were performed using the Brain Imaging Analysis Kit (BrainIAK v. 0.9.1; http://brainiak.org; Kumar et al., 2020, 2021) and Python 3.6, both in the subject’s native and functional space. A searchlight was then defined as a spherical cluster with a radius of 2 voxels (approximately 4.6 mm), encompassing 32 voxels around each target brain voxel. For each word, we averaged the whole-brain voxel activations across all instances into a single vector, resulting in 36 feature vectors representing 18 living and 18 non-living words. The dissimilarity between the average brain activations of different word pairs within each searchlight was quantified using the Pearson correlation distance, calculated with the Scipy library’s pdist function set to the ‘correlation’ metric (Virtanen et al., 2020). Similar to the models’ dissimilarity matrices, correlations between identical word pairs (e.g., dog - dog) and symmetrical pairs (e.g., dog - hammer and hammer - dog) were excluded. The neural data-derived RDM within each searchlight was then compared with the computational models’ RDM using the Spearman rank correlation (Carlin et al. 2011; Kriegeskorte et al., 2008). The Spearman ρ correlation coefficients obtained from this comparison were allocated to the central voxel of each searchlight, generating a whole-brain similarity map that depicted the correlations between brain activity patterns and computational model patterns.

To address the unknown null distribution in imaging data, a non-parametric statistical approach was used. To ensure a normal distribution of the correlation coefficients, the Spearman correlation values for each subject were Fisher-Z transformed, which stabilizes the variance and aids in subsequent statistical analyses. These Fisher-transformed correlation coefficient maps were then converted from the subject’s native space to the MNI-152 222 mm³ standard space using FSL. Within each group, a non-parametric one-sample t-test was conducted on participants’ MNI maps to identify brain regions where the Z-transformed correlation coefficients were significantly different from zero. For intergroup comparisons, two two-sample unpaired T contrasts were established: young group > older group, and older group > young group. The non-parametric test was carried out using FSL RANDOMISE, incorporating Threshold-Free Cluster Enhancement (TFCE cluster correction) and 10,000 random permutations (Nichols & Holmes, 2002; Smith & Nichols, 2009; Winkler et al., 2014). Variance smoothing, with a spatial FWHM Gaussian Kernel of 5 mm, was applied to enhance statistical power. Multiple comparisons were adjusted using Family-wise Error (FWE) correction. The corrected statistical image, produced by FSL RANDOMISE, was thresholded to highlight statistically significant clusters (p ≤ 0.05). The averaged Spearman correlation maps, showing significant correlations within groups and differences between groups, are presented below.

## Results

In this section, we present the results of our decoding analysis and RSA for both shallow and deep processing conditions. Initially, we assessed the decoding performance of a machine learning classifier within each condition independently, assessing how well the classifier can predict semantic categories based on brain activity. We then examined the generalization capabilities of the classifier, investigating how a decoder trained in one condition (e.g., shallow processing) performs when applied to another (e.g., deep processing) and vice versa. Additionally, our RSA results provide insights into the similarity between brain representations of experimental and those derived from a linguistic and computer vision model, across the entire brain. This comprehensive analysis provides insights into the neural encoding of semantic categories related to animacy under various cognitive processing loads and how these representations change as a function of aging.

### Within-condition decoding results

A leave-pair-words-out cross-validation approach was used within each condition (i.e., shallow, and deep processing) to assess decoding accuracy for unseen word pairs’ instances (corresponding to 1 living and 1 non-living concepts). The model was trained on a subset of 34 semantic concepts and tested on all possible pairs. This approach facilitated the assessment of decoding accuracy and the examination of how age, task condition, anatomical ROIs, and hemispheres influence the results using a mixed ANOVA analysis.

Decoding analyses within each level of processing (shallow, and deep processing) did not reveal any significant effect of aging or interactions with other factors. Hence, both age groups showed similar decoding patterns of semantic categories across the tested ROIs in both shallow and deep conditions, and across both hemispheres. However, we observed that decoding performance was better in the left hemisphere compared to the right hemisphere. Furthermore, the overall decoding accuracies were significantly greater in the deep processing condition than in the shallow processing condition across both hemispheres in line with the observation reported in a recent study involving young adults (Soto et al., 2020; see also the Supplementary results for a detailed account of the statistics unrelated to the effects of age). Figure 3 provides an overview of the decoding results across ROIs, experimental conditions, hemispheres, and age groups.

### Cross-condition decoding results

We then examined the impact of aging on the specificity and generalization of brain representations of semantic categories across the different levels of processing. We utilized a leave-pair-words-out cross-validation method, where a model was trained using data from 34 out of 36 semantic concepts in one task condition (e.g., deep), and then assessed on all potential pairs of concepts that were excluded (1 living concept, 1 non-living concept) in the other task condition (e.g., shallow). This resulted in two cross-condition directions: shallow-to-deep and deep-to-shallow. Figure 4 illustrates the cross-condition decoding results across various ROIs, cross-condition direction, hemispheres, and age groups. Decoding accuracies were compared utilizing a mixed ANOVA across ROIs, cross-condition directions, hemispheres, and age groups.

Similar to the within-condition decoding results, the cross-condition decoding analysis revealed significant main effects and also interaction effects that were unrelated to aging. These results are presented in the Supplementary results. Here, we report the most critical results regarding the effects of aging on cross-condition decoding.

The ANOVA revealed a significant main effect of age [F(1, 54) = 4.315, p = .043, 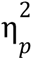 = .074], indicating that cross-condition decoding was higher in the older group (Mean = .524, SE = .004) compared to the younger group (Mean = .513, SE = .004). We also observed an interaction effect of age and ROIs [F(12.650, 683.082) = 2.269, p = .007, 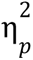 = .04]. To unravel the source of this interaction, post-hoc pairwise comparisons were conducted, averaging over hemisphere and cross-condition direction levels. Within each ROI, unpaired two-tailed t-tests, with p-values corrected for multiple tests using the FDR method, were performed to compare the two age groups. Notably, a significant effect was found in the middle temporal lobe, with cross-condition decoding being significantly higher in the older group (Mean = .542, SE = 6.4e-3), compared to the young group (Mean = .514, SE = 5.3e-3) [t(54) = 3.300, p = .026].

We further observed a significant interaction effect among age, ROIs, and hemisphere [F(11.562, 624.340) = 1.862, p = .038, 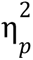 = .033] indicating that the main effect of aging on cross-condition decoding depended on the hemisphere and ROI. To further investigate this interaction, two supplementary mixed-effects ANOVA analyses were conducted independently in the left and right hemispheres, utilizing a 15 (ROIs), 2 (cross-condition direction: shallow-to-deep vs. deep-to-shallow), 2 (age: older group vs. young group) design with Huynh-Feldt corrected DOF.

The supplementary ANOVA of the right hemisphere showed no significant main effect of age or interactions with other factors. However, in the left hemisphere supplementary ANOVA, a main effect of age was observed (F(1, 54) = 5.273, p = .026, 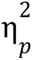 = .089). Specifically, the older group (Mean = .533, SE = .004) exhibited larger cross-condition decoding compared to the younger group (Mean = .519, SE = .004). Additionally, an age by ROIs interaction was found (F(12.056, 651.035) = 2.962, p < .001, 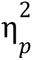 = .052). To explore this interaction effect, post-hoc pairwise comparisons were performed, collapsing across cross-condition decoding direction levels. Within each ROI in the left hemisphere, unpaired two-tailed t-tests with p-values corrected using the FDR method were conducted to compare the two age groups. Notably, significant differences were observed in the left inferior parietal lobe [Older: Mean = .566, SE = .010; Young: Mean = .528, SE = 6.4e-3; t(54) = 3.092, p = .024] and the left middle temporal lobe [Older: Mean = .566, SE = 9.9e-3; Young: Mean = .518, SE = 7.5e-3; t(54) = 3.780, p = .006] (Figure 5).

**Figure 5.**
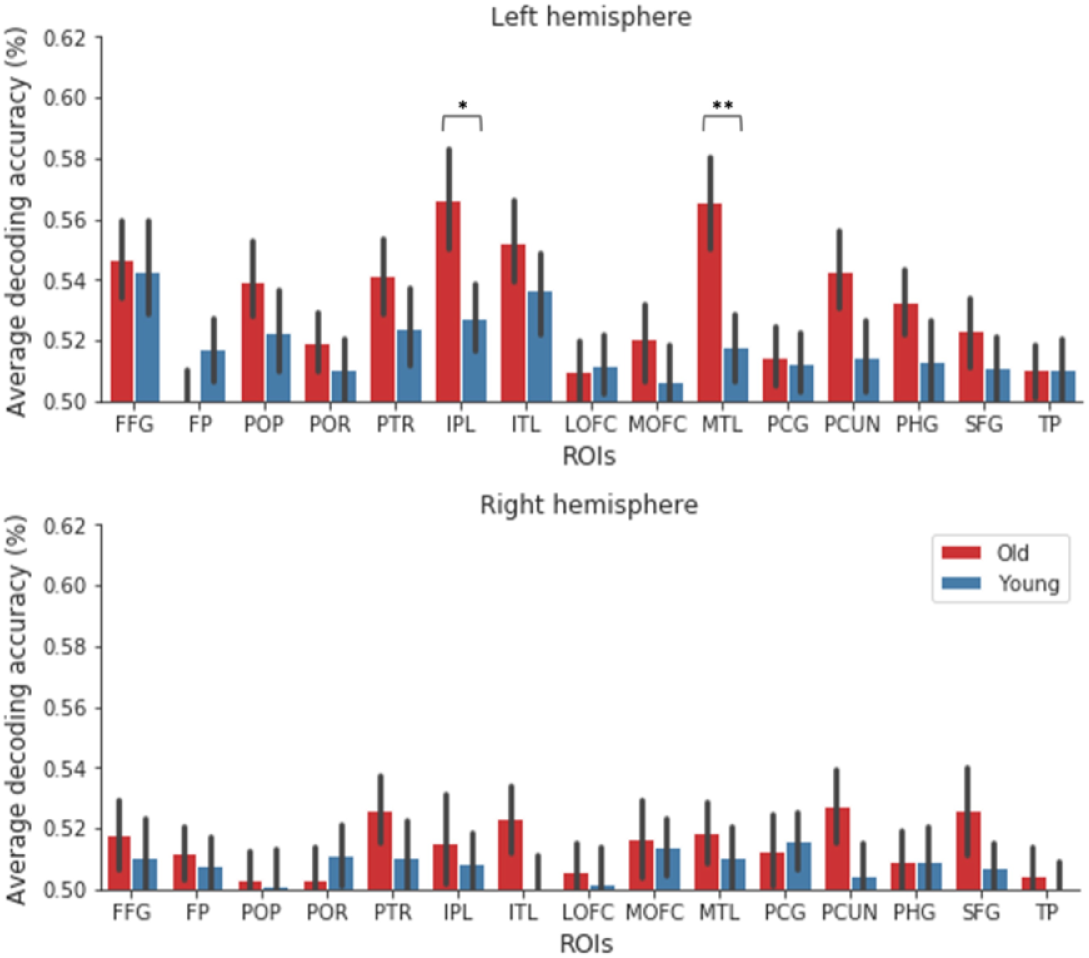
Cross-condition decoding accuracy averaging over cross-condition direction levels (i.e., deep-to-shallow, and shallow-to-deep) by ROI, hemisphere, and age-group. The error bars represent a 95% confidence interval. * p < .05, ** p < .01.

### RSA results

We used searchlight RSA to compare how the brain represents semantic categories of animacy to the computational representations provided by fine-tuned language and computer vision models (specifically, Word2Vec and VGG-19). Additionally, we examined how these representations differ across age groups.

To evaluate the significance of the correlations between each model and the brain representational dissimilarity matrices, we began by conducting one-sample permutation t-tests on the searchlight RSA maps for each age group separately. This approach aimed to identify brain clusters where the correlations were significantly greater than zero, applying whole-brain correction and a threshold of p < 0.05 with 10,000 permutations.

In the shallow processing condition, significant correlation clusters were found exclusively in the older group. For both the computer vision and word embedding models, these clusters were primarily located in the left middle temporal gyrus. In contrast, in the deep processing condition, significant correlation clusters were found in both the young and older groups. In keeping with the decoding analyses conducted within each depth of processing condition, the strongest correlations between neural data and latent features from VGG-19 and Word2Vec were observed in the left hemisphere. In both models, the correlations were particularly strong during the deep condition, spanning regions across the ventral visual pathway (including the occipitotemporal cortex, middle temporal gyrus, and inferior temporal gyrus), the dorsal visual pathway (such as the angular gyrus, supramarginal gyrus, superior parietal lobule, postcentral gyrus, parietal operculum cortex, and precentral gyrus), and frontal regions like the middle frontal gyrus, inferior frontal gyrus, and frontal pole.

We then compared the RSA maps across age groups, revealing significant differences in the deep processing condition. Specifically, the brain RDMs of the younger group showed higher correlations with the computational models’ RDMs compared to the older group, particularly in the bilateral postcentral gyrus and surrounding regions. These differences were most pronounced in the left hemisphere and were consistent for both the VGG-19 and Word2Vec models (Figure 6; model RDMs are presented in Figures 7A and 7B). No significant differences were observed between age groups in the shallow processing condition.

**Figure 6.**
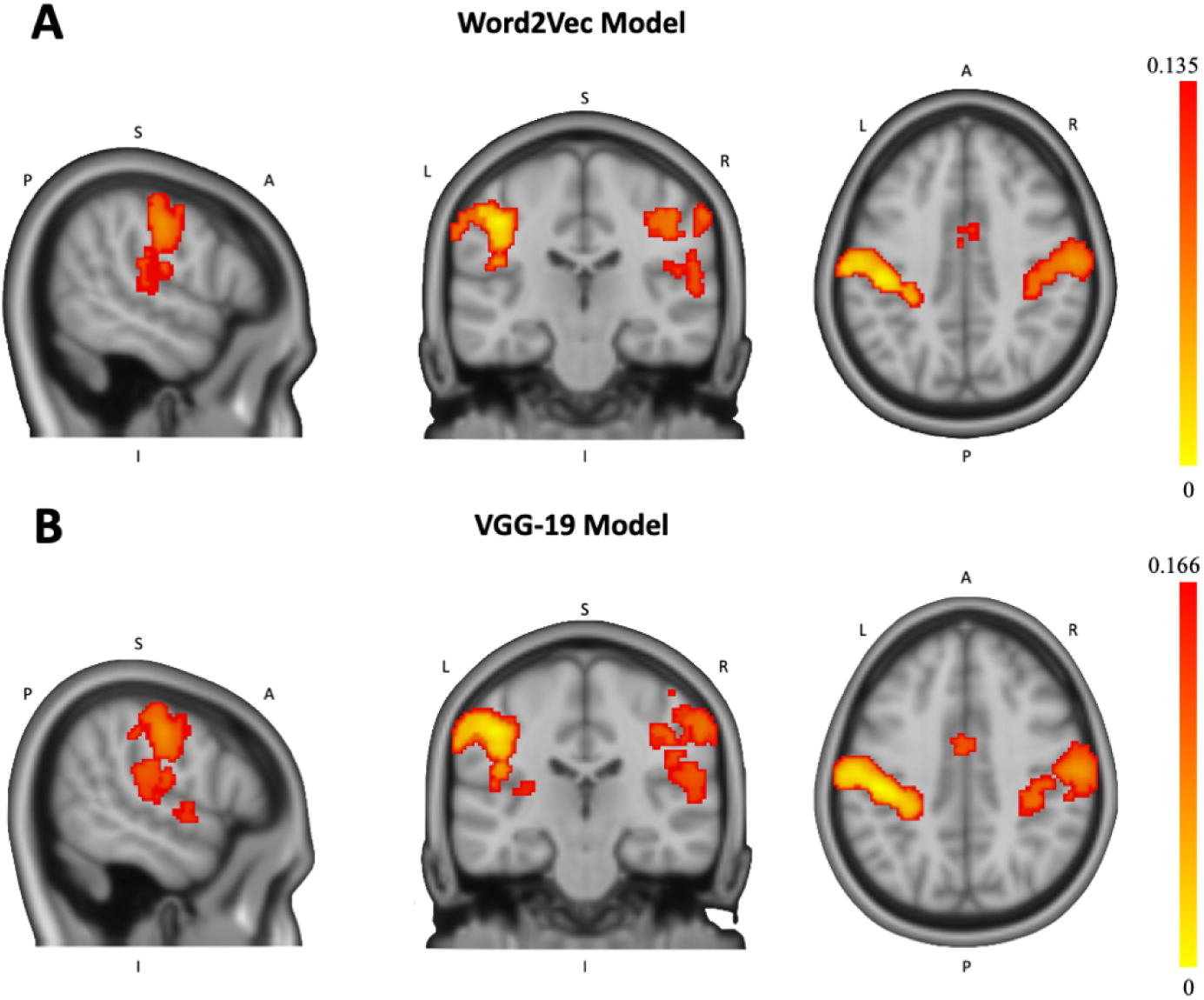
Deep processing condition. Searchlight RSA correlation differences between age groups in the young > older direction using the word embedding model (A) Word2Vec, and the computer vision model (B) VGG-19.

**Figure 7.**
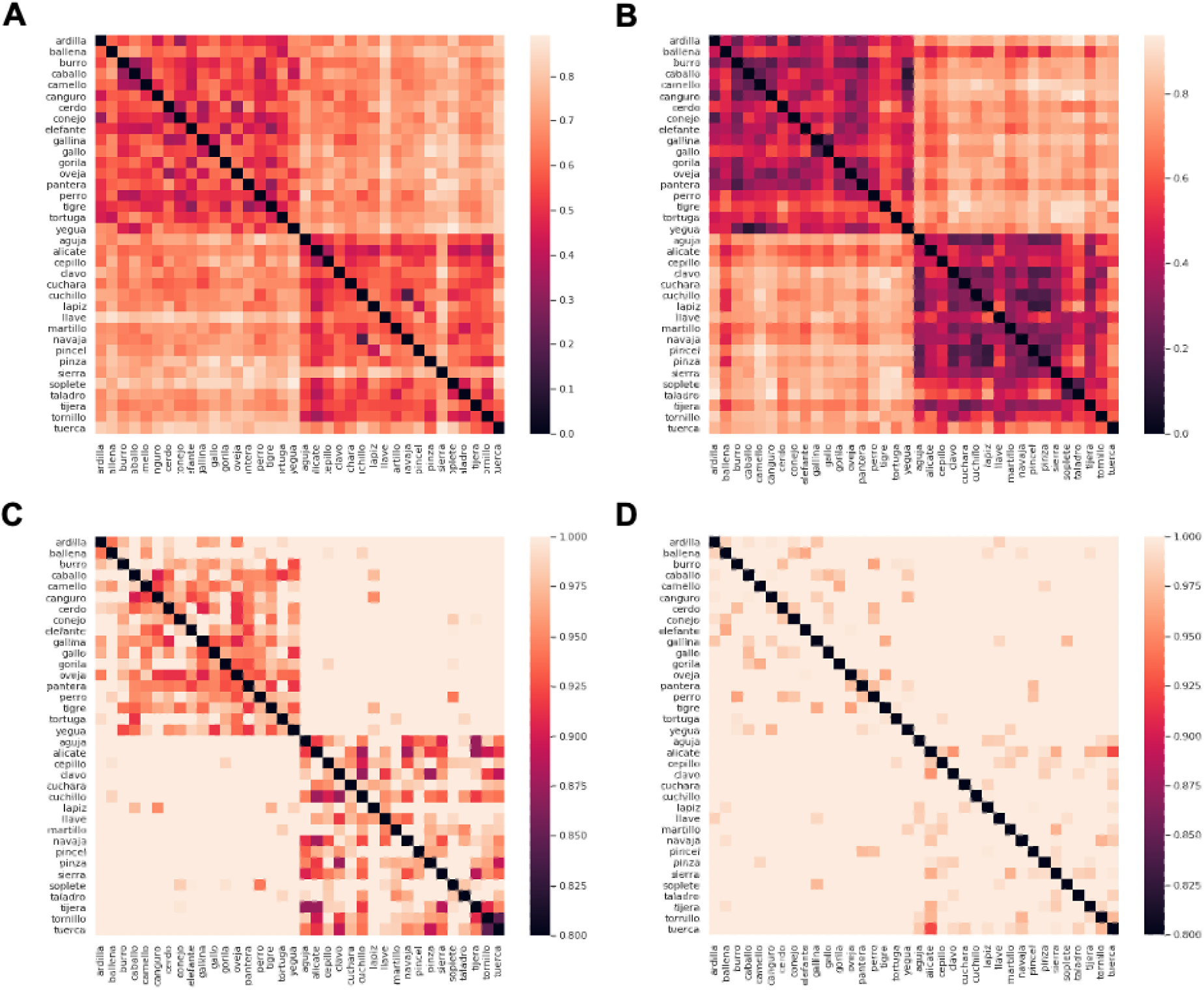
(A) Representational dissimilarity matrix (RDM) of the fine-tuned Word2Vec model embeddings. (B) RDM of the fine-tuned VGG-19 model embeddings. (C) Average RDM of brain activation patterns from the searchlight analysis in the postcentral gyrus cluster for the young group. (D) Average RDM of brain activation patterns from the searchlight analysis in the postcentral gyrus cluster for the older group. The RDMs represent concepts for each living and non-living category. In the figures, the top-left squares show the dissimilarity among concepts within the living condition. The bottom-left squares show the dissimilarity among concepts across both the living and non-living categories. Lastly, the bottom-right squares show the dissimilarity among concepts within the non-living condition. A darker shade of red indicates a higher degree of representational similarity between concepts (p), while a lighter shade of red suggests greater dissimilarity (1- ρ).

However, further examination is needed to precisely understand the nature of the age differences at the representational level. It remains unclear whether the neural representation of different animacy categories in the postcentral gyrus cluster become more similar with aging, or whether aging also impacts the similarity between neural representations of different items within each category.

Data from all participants were transformed into MNI space to extract information from the significant postcentral gyrus cluster. For each participant, we calculated the average Pearson dissimilarity scores (1 - ρ) for all pairs of concepts within each living and non-living category, as well as between categories. The correlation distance values for each subject were subjected to the Fisher-Z transformation for normality.

A mixed ANOVA with category as a within-subjects factor (living, non-living, average within-categories, and between categories concepts dissimilarity) and age group as between-subjects factor (young vs. older adults) was conducted. Significant main effects were found for both the category and group factors, as well as a significant interaction effect between these factors [F(3, 162) = 12.051, p < .001, 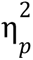 = .182].

Post-hoc t-tests with Holm-Bonferroni correction for multiple comparisons revealed that older adults exhibited greater dissimilarity among concepts within the living [Mean Difference = .041, SE = .011, t(55) = 3.736, p = .004], within the non-living category [Mean Difference = .035, SE = .011, t(55) = 3.250, p = .016], and also for the average dissimilarity of both categories (i.e., average within-category dissimilarity) [Mean Difference = 0.038, SE = .011, t(55) = 3.493, p = .009]. However, the older group showed higher similarity between the living and non-living concepts compared to the young group [Mean Difference = -.036, SE = .011, t(55) = -3.356, p = .012]. These results are depicted in Figure 7C and 7D, and Figure 8A.

**Figure 8.**
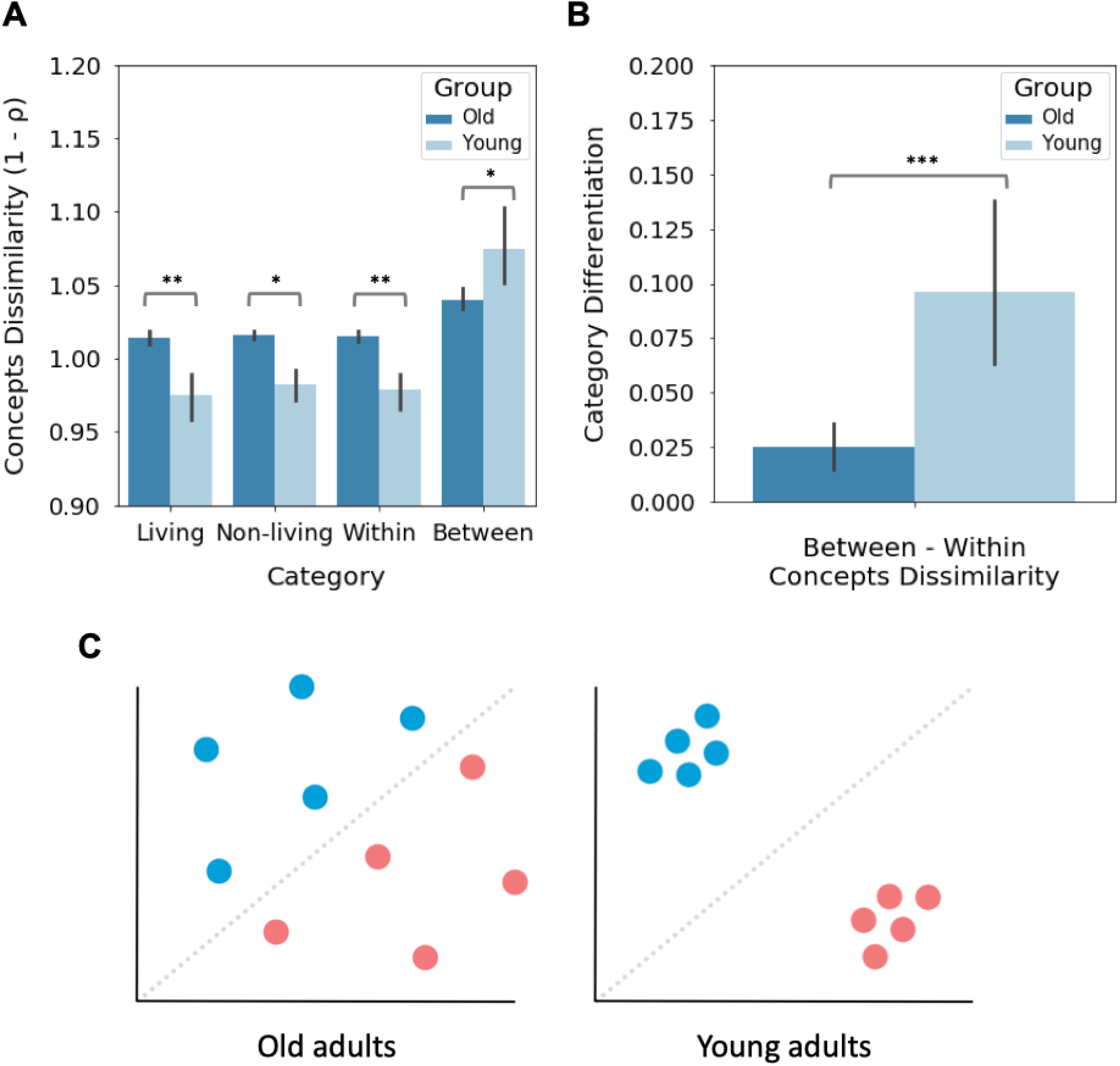
(A) Comparison of average dissimilarity in conceptual representation across living, non-living, and combined categories, as well as between different category concepts. (B) Quantification of category distinctiveness within each group by computing the difference between dissimilarity among concepts from different categories and dissimilarity among concepts within the same category. The error bars represent the 95% confidence interval, adjusted using the Cousineau-Morey method (Cousineau, 2005; Morey, 2008). * p < .05, ** p < .01, *** p < .001. (C) The figure illustrates the hypothetical representational structure derived from RSA results for each group, showing the dissimilarity of concepts within and between categories. Blue dots correspond to concept representations from one specific category (e.g., living), while red dots represent concept representations from the other category (e.g., non-living). In older adults, there is increased dissimilarity in conceptual representations within each category and increased similarity between concepts from different categories. Conversely, young adults show the opposite pattern.

Finally, we compared the neural pattern of dissimilarity between categories across groups by subtracting the within-categories concept dissimilarity from the between-categories dissimilarity for each group (Carp, Park, Hebrank, et al., 2011; Deng et al., 2021; Haxby et al., 2001). An independent sample t-test demonstrated that the older group [Mean = .026, SE = .006] exhibited enhanced category similarity in neural patterns compared to the young group [Mean = .100, SE = .021] [t(54) = -3.492, p < .001] (Figure 8B).

## Discussion

The present fMRI study used a comprehensive approach including pattern classification, and searchlight RSA analyses based on computational models, with the goal of defining the impact of aging on the brain representation of living and non-living semantic categories and how this varies across different levels of processing associated with the task.

Although there were no differences in decoding accuracy within individual tasks between the young and older groups, we observed that in the cross-condition generalization analyses, decoding accuracy was significantly higher in older adults compared to the young group. These analyses can be interpreted in light of the neural dedifferentiation hypothesis of aging (Koen & Rugg, 2019), according to which aging leads to a reduction in the distinctiveness of brain activity patterns. Previous work used MVPA methods to assess age-related changes in neural representations in visual processing (Carp, Park, Polk, et al., 2011), motor responses (Carp, Park, Hebrank, et al., 2011), episodic memory (Abdulrahman et al., 2017) and working memory (Carp et al., 2010). These investigations have shown that, compared to younger individuals, older adults tend to display less specific brain activity patterns for task relevant representations in various domains (e.g., for visual categories such as faces and houses; Park et al., 2010). Similarly, our cross-condition decoding results indicate that the brain representations of semantic categories become more similar with aging across different task conditions which differ in the depth of processing, specifically in the left inferior parietal and middle temporal lobes that are known to be implicated in the storage of lexico-semantic information during language processing (Binder & Desai, 2011). These results indicate that aging leads to generalized neural representations of semantic knowledge.

Furthermore, our model-based searchlight RSA indicated that the brain representations related to animacy in the young group closely resembled the fine-grained distinction of living and non-living concepts according to the language model and computer vision model. However, older participants displayed a reduction in pattern similarity relative to the models’ RDMs in the superior parietal during the deep processing condition. A recent fMRI study by Deng et al. (2021) examined age-related differences in visuo-perceptual and mnemonic representations of complex scenes using RSA. They compared representations from the initial and penultimate hidden layers of a computer vision model (VGG-16; Simonyan & Zisserman, 2014) with the participants’ brain representations of the images. The findings showed that in older adults, brain activity patterns in the early visual cortex were less aligned with the image model based on the initial layer, which encodes basic visual features such as edges, colors, and textures, compared to young adults, supporting the neural dedifferentiation hypothesis. In contrast, they found a stronger correlation between brain activity patterns in the ventral anterior temporal lobe and the higher-order categorical layer of the computer vision model—which processes abstract category information—in older adults compared to young adults. This finding, which aligns with results from Naspi et al. (2021, 2022), was interpreted as neural hyperdifferentiation, indicating a greater distinctiveness of higher-order neural representations. In our study, the searchlight RSA and cross-condition decoding results support the presence of neural dedifferentiation in the semantic domain; however, we found no evidence of neural hyperdifferentiation. The discrepancies between our RSA results and previous studies might be attributed to variations in the cognitive processes and stimuli targeted by each study. In the previous studies, participants were specifically tasked with encoding and visually recalling images of complex scenes, as well as living and non-living objects, emphasizing visual processing and episodic memory. Our study extends the work of Deng et al. and Naspi et al. by exploring more general semantic processes. It allows participants to freely engage in multimodal mental simulations of concepts, encouraging the integration of information from various sensory modalities, such as visual, somatosensory, auditory, and memory. The postcentral gyrus, serving as the primary somatosensory cortex, plays a crucial role in processing and integrating sensory information related to touch, proprioception and generating motor responses (Kropf et al., 2018; Longo et al., 2010). As individuals age, changes in body perception and action, such as tactile perception, balance, interoception, proprioception, manual motor control, and walking, are commonly observed (see Kuehn et al., 2018 for a review). These age-related declines in physical and cognitive functions can impact the integration of sensory-motor information in the postcentral gyrus associated with simulation of living and non-living word concepts. This perspective is consistent with previous research on neural dedifferentiation, which found that the distinctiveness of neural representations of motor actions was also significantly higher in young adults compared to older adults in bilateral neighboring regions to the postcentral gyrus, including the primary motor cortex and presupplementary motor area (Carp, Park, Hebrank, et al., 2011). Intriguingly, the RSA analyses also revealed an age-related reduction in neural pattern similarity (i.e., increased distinctiveness) of item-based conceptual representations within the living and non-living categories. The formulation of neural dedifferentiation in previous studies (e.g., Carp, Park, Hebrank, et al., 2011; Deng et al., 2021) focused primarily on the comparison of neural pattern similarity within and between conditions, as proposed by Haxby et al. (2001). Our study builds upon this conceptualization but, critically, highlights the significance of addressing neural pattern similarity at different levels of semantic analysis, both at the lowest dimensional level (conceptual, item-level) and the higher-dimensional level (category) of analysis. Moreover the juxtaposition of decoding and RSA results in our study, underscores the importance of employing diverse analytical methods when investigating semantic neural dedifferentiation. While decoding analyses employing SVM models predominantly aim to maximize the margin between categories, RSA takes into account the fine-grained pattern differences across all concept pairs, offering a more comprehensive view of the representational space.

When considering potential limitations of our study, one could argue that our results, particularly the increased cross-condition generalization observed in older adults, might stem from carry-over effects between conditions. Previous studies have shown that older adults tend to exhibit reduced attentional control and executive functions (Paxton et al., 2008; Verhaeghen et al., 2003; Verhaeghen & Cerella, 2002; Wasylyshyn et al., 2011). Consequently, it might be argued that older adults unintentionally engaged in mental simulation during the covert reading task, and vice versa, due to task-switching costs across the different fMRI runs. However, this explanation seems unlikely. First, it should be noted that, although we included catch trials, due to the covert nature of our tasks, we could only monitor the quality of participants’ performance by means of neural decoding. However, we implemented several strategies to ensure participants engaged in the tasks in an active manner. For example, participants received thorough instructions and engaged in practice sessions before scanning to familiarize themselves with the task requirements. Following each fMRI run, regular check-ins and debriefings were held to respond to any queries from participants and to evaluate their engagement. Furthermore, the inclusion of catch trials should increase participants’ compliance with the instructions and the data indeed showed this to be the case. Second, if carry-over effects were present, they would have been noticeable in the shallow processing condition due to switching costs from the previous deep processing run. This scenario should have resulted in significant decoding accuracies among older adults in the shallow condition during within-condition decoding analyses. However, this pattern of results was not evident. Older adults did not exhibit significant differences in within-condition analyses when compared to young adults, in both shallow and deep processing conditions, indicating that both age groups likely utilized similar mental strategies during task performance. Moreover, our analysis of catch trials did not reveal any significant differences between age groups, suggesting similar levels of attention and engagement in the specific conditions of our experiment. An additional potential concern relates to the fact that the different concepts reappeared across runs and the possibility of age-related differences in neural repetition suppression (Bergerbest at al., 2009). It may be argued that these age-related differences in neural repetition suppression could impact on the decoding results. However, the absence of age-related differences in decoding accuracy within the shallow or deep processing conditions indicates that the age-related differences observed in the cross-condition analyses are unlikely to be mediated by differences in neural repetition suppression effects.

To conclude, our study highlights the dynamic nature of semantic representations in the aging brain and suggests that healthy aging is characterized by complex changes in neural patterns, rather than by a uniform loss of neural pattern specificity in semantic representations. Deng et al. (2021) and Naspi et al. (2022) observed that neural dedifferentiation and hyperdifferentiation can co-exist in the representation of visual categories, and that neural hyperdifferentiation may be a compensatory mechanism to counteract the diminished quality of sensory representations in healthy aging. However, Deng et al and Naspi did not address how these processes operate regarding general semantic knowledge at the interface between language and cognition. The present study provides novel insights by highlighting how neural representational dedifferentiation (i.e. increased similarity) and hyperdifferentiation (i.e. increased specialization) in aging operate across the different levels of semantic analysis, from basic conceptual or item level representations to broader, higher-dimensional category levels. This emphasis is particularly crucial in the semantic domain because concepts serve as meaningful units and contribute to the compositionality of language. We further propose that neural dedifferentiation in the semantic domain may also represent a beneficial compensatory mechanism (Seider et al., 2021), helping to maintain semantic processing in aging by allowing for a more efficient coding scheme, both compressed and sparse, to make better use of limited neural resources in healthy aging.

## Acknowledgments

D.S. acknowledges support from the Basque Government through the BERC 2022-2025 program and from grant PI_2017_1_0025, and also from the Spanish State Research Agency, through the ‘Severo Ochoa’ Programme for Centres/Units of Excellence in R & D (CEX2020-001010-S).

## Supplementary materials

**Table S1.**
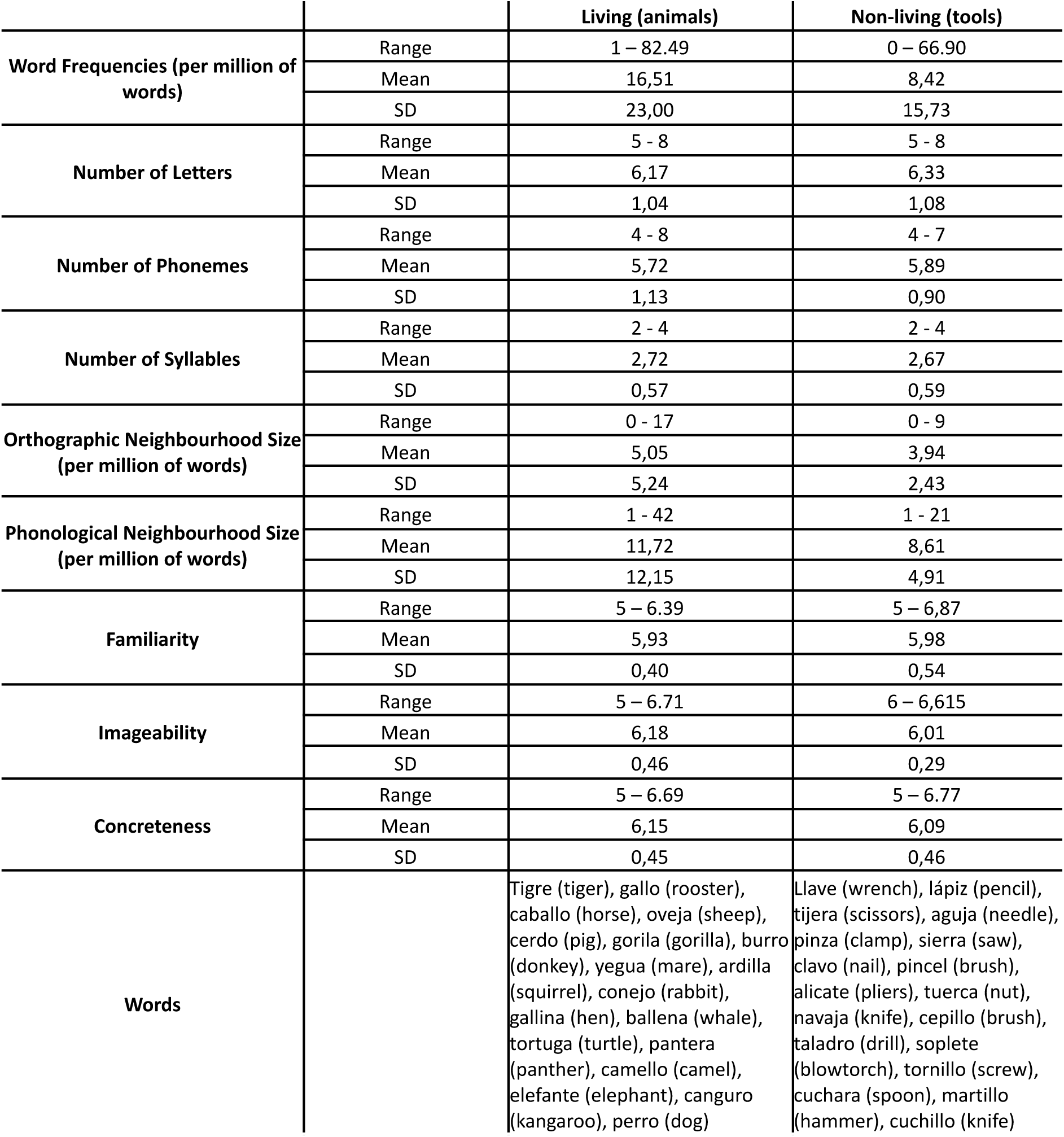
Summary of living and non-living Spanish words’ properties gathered from EsPal database (bcbl.eu/databases/espal/; Duchon et al., 2013). There were a few words (12) for which some familiarity, imageability, and concreteness ratings were not found on this database. Reported range, mean and SD of ratings for familiarity, imageability and concreteness were calculated only for the words for which it was found. Independent samples t-tests, adjusted for multiple comparisons, were conducted to compare the properties of living and non-living words. No significant differences were observed between the word categories for any of the properties.

**Figure S1.**
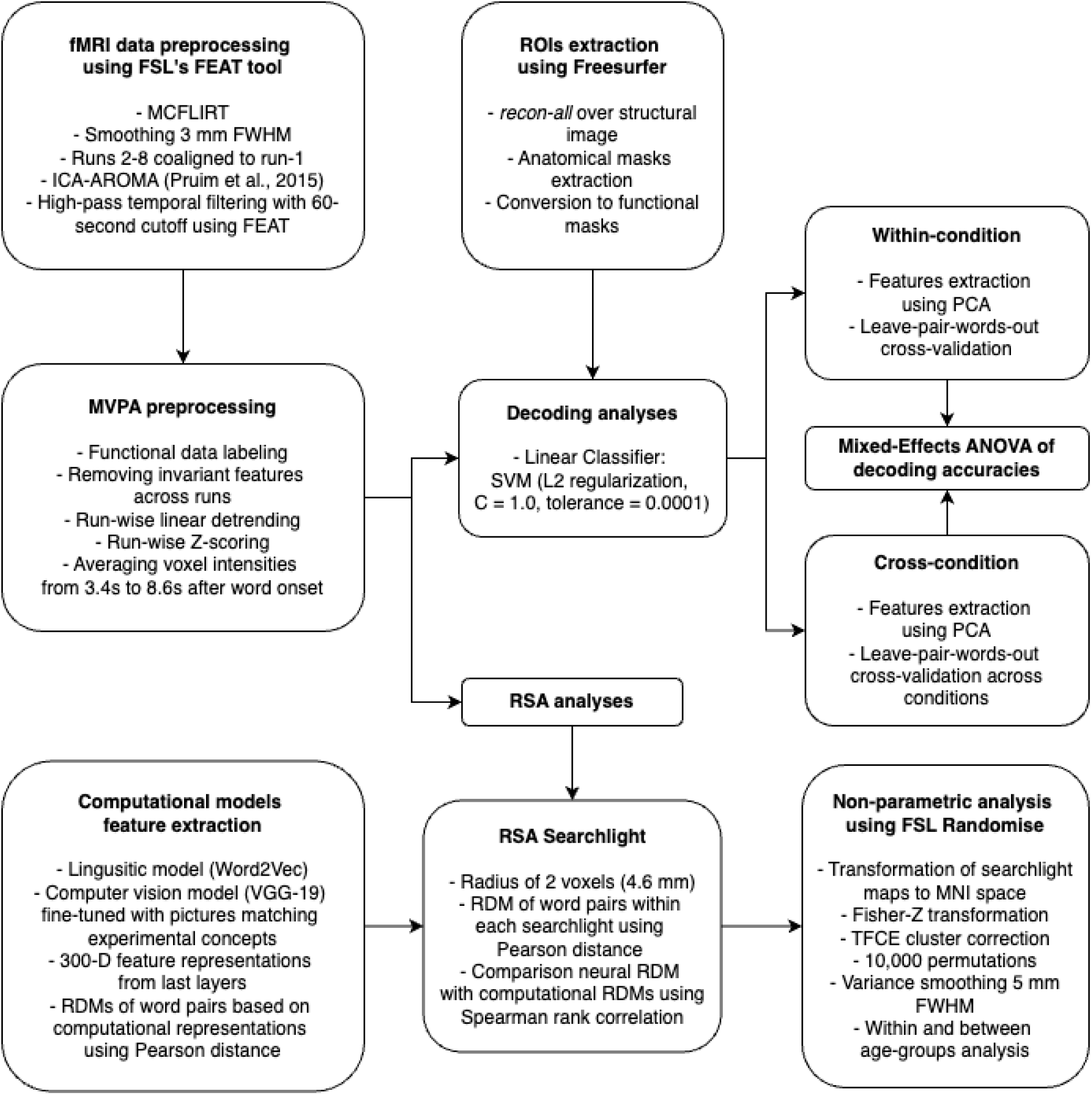
Comprehensive overview of the MVPA pipeline.

### Word embedding model

In our study, we employed the Word2Vec word embedding model, chosen for its established performance in prior studies (Fernandino et al., 2022; Martin et al., 2018; Wang et al., 2018) and the availability of a pre-trained version trained on the Spanish Billion Word Corpus (Cardellino, 2016). This model, accessible from the GitHub repository at github.com/dccuchile/spanish-word-embeddings, captures both the semantic and syntactic properties of words as dense numerical vectors (embeddings). Words with similar meanings have similar embedding characteristics. We adhered to common practices in natural language processing by extracting 300-dimensional vector representations for each of the 36 words from the final layer of the pre-trained Word2Vec model. To standardize these feature representations, we subtracted the mean and divided by the standard deviation of each word’s vector. The representational dissimilarity between word pairs was calculated using the Pearson correlation distance, employing the Scipy library’s pdist function with ‘correlation’ specified as the distance metric (Virtanen et al., 2020). To maintain analytical robustness, self-correlations (e.g., dog - dog) and symmetrical pairs (e.g., dog - hammer and hammer - dog) were excluded from the calculations. Figure 7A showcases a visualization of the RDM derived from the pairwise correlations of the embeddings, illustrating how each word’s representation diverges from others in the set.

### Computer vision model

We selected the VGG-19 model (Simonyan & Zisserman, 2014), a pre-trained computer vision model known for its relatively shallower architecture of 26 layers, as a robust baseline in previous research (Deng et al., 2021; He et al., 2016; Naspi et al., 2022; Ren et al., 2015; Russakovsky et al., 2015). This convolutional neural network is specifically designed to map images to dense feature vectors and generate category probability estimates, making it a suitable choice for our study.

The model, pre-trained on the ImageNet dataset (imagenet.org), was sourced from the Keras Python Library (Chollet, 2015). It’s important to acknowledge that the dimensionalities of computer vision models and word embedding models typically differ. While the VGG-19 produces 512-dimensional (512-D) feature vectors for images, we modified the computer vision model to align with the 300-dimensional (300-D) vectors of the word embedding model to ensure comparability. This was done by adding a new layer to the pre-trained model’s structure before fine-tuning it (Mesnil et al., 2012; Yosinski et al., 2014). During its pre-training phase, the VGG-19 model was initially trained to identify the category of an input image from the ImageNet dataset, which comprises a thousand different categories. This pre-training imbued the model with valuable insights that were further refined for application to our specific image dataset and classification needs. To optimize the VGG-19 for our specific word-image dataset, we engaged in a fine-tuning process using the extensive dataset from the Caltech 101 database (Fei-Fei et al., 2007), which features 101 different object categories. This dataset offers a broad spectrum of images representing a wide range of visual concepts, accessible via the URL: data.caltech.edu/records/mzrjq-6wc02. The image categories in the Caltech 101 database include between 45 and 400 images each. Before initiating fine-tuning, we excluded six categories that did not align semantically with the living and non-living categories central to our experiment. These were: “BACKGROUND google”, “Faces”, “Faces easy”, “Brain”, “Stop sign”, and “Trilobite”. To maintain a balanced sample distribution across categories, we employed a stratified sampling method, selecting 30 images randomly from each category. Of these, 24 images were used for training, and 6 for testing. All images were normalized individually according to the specific preprocessing guidelines detailed in the Keras documentation and resized to dimensions of 128 x 128 pixels. To further tailor the model to our classification task, we augmented the architecture by adding a new classification layer designed to handle 96 categories on top of the existing 300-D layer. This adaptation ensured the model was well-equipped for our specific classification requirements.

The fine-tuning process involved several steps to adapt the VGG-19 model to our experimental settings. Initially, we locked the convolution pooling blocks of the pre-trained model, ensuring that during the training and validation phases, only the weights of the newly added layers were subject to adjustments. To enhance the model’s performance, we applied various augmentation techniques to the image sets, including rotation, shifting, zooming, and flipping. For the newly incorporated 300-D layer, we employed the Self-Normalizing Neural Networks (SNNs) activation function (Klambauer et al., 2017), and initialized the weights using the LeCun normal initialization method (LeCun et al., 2012) as outlined in the TensorFlow documentation (Abadi et al., 2016). The optimization of the model was handled by the Adam optimizer, set at a learning rate of 0.0001, and the categorical cross-entropy function was chosen as the loss metric (Kingma & Ba, 2014). No additional regularization methods were used. The model was trained for up to 3000 epochs with a batch size of 16 images per back-propagation step. Training was halted if the model’s performance on the validation set failed to improve for five consecutive batches.

Following the fine-tuning of the VGG-19 model, it was utilized to extract abstract representations of image referents corresponding to the experimental words. For each of the 18 living and 18 non-living words, a set of 10 images depicting the objects or entities represented by these words was gathered from the Internet. These images were cropped to center the object against a white background and were normalized according to the specifications of the selected model. Subsequently, these prepared images served as input data for the fine-tuned computer vision model. The penultimate layer, which is the newly added 300-D layer, produced the output vectors representing the features of each image referent. To achieve a shared representation, the vector outputs for all 10 images associated with each word were averaged. Similar to the word embedding models, these 300-D feature representations for the images were standardized, and their pairwise correlations were calculated to construct the RDM, as shown in Figure 7B. This process ensured a consistent and comparative analysis of the image-based features alongside the linguistic data.

## Supplementary results

### Within and cross-condition decoding analyses

We compared the decoding accuracies obtained from within and cross-condition generalization analyses using ANOVA. These analyses involved investigating the effects of various factors and their corresponding levels. Specifically, for within-condition analyses, the factors included 15 ROIs, 2 conditions (shallow vs. deep), 2 hemispheres (left vs. right), and 2 age groups (young vs. older). In the case of cross-condition analyses, the factors consisted of 15 ROIs, 2 cross-condition directions (shallow-to-deep, deep-to-shallow), 2 hemispheres (left vs. right), and 2 age groups (young vs. older). Although our primary focus revolved around exploring the impact of age as a factor within and across conditions when decoding multi-voxel patterns for unseen word pairs, this section provides a detailed exploration and discussion of the ANOVA results pertaining to additional factors.

#### Increased decoding of semantic categories in the left hemisphere in both within condition and cross-condition analyses

In the within-condition decoding analysis, we observed a significant main effect of the hemisphere factor [F(1, 54) = 42.905, p < .001, 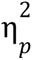 = .443]. Decoding accuracies were significantly higher in the left hemisphere (Mean = .536, SE = .005) compared to the right hemisphere (Mean = .524, SE = .005) (Figure S2). Additionally, we found interactions between the hemisphere factor and other variables: hemisphere * ROIs [F(11.856, 640.216) = 7.057, p < .001, 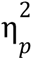 = .116], hemisphere * condition [F(1, 54) = 15.975, p < .001, 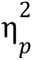 = .228], and hemisphere * condition * ROIs [F(11.927, 644.039) = 2.342, p = .006, 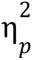 = .042]. In the subsequent paragraph, our primary objective is to delve deeper into the interaction between hemisphere, condition, and ROIs. Our aim is to examine the variations in decoding accuracies between hemispheres within both the shallow and deep conditions for each specific ROI.

We conducted two additional mixed-effects ANOVA analyses, one in the shallow condition and another in the deep condition using 15 ROIs, 2 hemispheres, and 2 age groups as factors. The DOF for these ANOVAs were corrected using Huynh-Feldt adjustment. In both the shallow condition [F(1,54) = 7.240, p = .009, 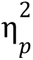 = .118] and the deep condition [F(1, 54) = 45.795, p < .001, 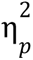 = .459], we found a significant main effect of the hemisphere. This indicates that overall decoding accuracies were higher in the left hemisphere [Shallow: Mean = .513, SE = .006; Deep: Mean = .560, SE = .008] compared to the right hemisphere [Shallow: Mean = .507, SE = .006; Deep: Mean = .541, SE = .008]. Furthermore, in the deep condition, we also observed a significant hemisphere * ROIs interaction [F(12.099, 653.331) = 7.351, p < .001, 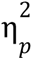 = .120]. To analyze this interaction effect, we conducted post-hoc pairwise comparisons, averaging over the age levels. Within each ROI, paired two-tailed t-tests were performed to compare decoding accuracies between the left and right hemispheres in the deep condition, with p-values corrected using FDR for multiple comparisons. In the deep condition, decoding accuracies were significantly higher in the left hemisphere compared to the right hemisphere in the following ROIs: fusiform gyrus [Left: Mean = .578, SE = .013; Right: Mean = .557, SE = .012, t(54) = 2.813, p = .017], inferior parietal lobe [Left: Mean = .607, SE = .013; Right: Mean = .570, SE = .011, t(54) = 5.193, p < .001], middle temporal lobe [Left: Mean = .592, SE = .011; Right: Mean = .556, SE = .011, t(54) = 5.065, p < .001], superior frontal gyrus [Left: Mean = .578, SE = .011; Right: Mean = .552, SE = .010, t(54) = 4.986, p < .001], inferior temporal lobe [Left: Mean = .613, SE = .013; Right: Mean = .550, SE = .012, t(54) = 8.40, p < .001], and pars opercularis [Left: Mean = .577, SE = .012; Right: Mean = .545, SE = 9.2e-3, t(54) = 3.980, p < .001] (Figure 3).

In the cross-condition decoding approach, we observed a significant main effect of the hemisphere factor [F(1, 54) = 33.411, p < .001, 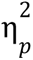 = .382]. Overall, cross-condition decoding accuracies were higher in the left hemisphere (Mean = .526, SE = .003) compared to the right hemisphere (Mean = .511, SE = .003) (Figure S3). Additionally, we identified two interactions: hemisphere * ROIs [F(11.562, 624.340) = 4.921, p < .001, 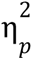 = .084] and hemisphere * cross-condition direction [F(1, 54) = 6.919, p < .001, 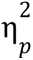 = .114].

First, we examined the hemisphere * ROIs interaction effect by conducting post-hoc pairwise comparisons, averaging across the levels of age and cross-condition direction. Paired two-tailed t-tests, with p-values corrected using the FDR for multiple comparisons, were performed within each ROI to compare decoding accuracies between the left and right hemispheres. We found significantly larger cross-condition decoding accuracies in the left hemisphere when compared to the right hemisphere in the fusiform gyrus [Left: Mean = .545 SE = 6.4e-3, Right: Mean = .514 SE = 5.4e-3, t(54) = 4.855, p < .001], inferior parietal lobe [Left: Mean = .548 SE = 6.7e-3, Right: Mean = .512 SE = 6.1e-3, t(54) = 4.750, p < .001], middle temporal lobe [Left: Mean = .543 SE = 7.0e-3, Right: Mean = .515. SE = 4.5e-3, t(54) = 3.742, p < .001], inferior temporal lobe [Left: Mean = .545 SE = 5.8e-3, Right: Mean = .512 SE = 5.4e-3, t(54) = 4.589, p < .001], pars opercularis [Left: Mean = .531 SE = 6.1e-3, Right: Mean = .502 SE = 4.8e-3, t(54) = 3.978, p < .001].

Next, we analyzed the hemisphere by cross-condition direction interaction. Post-hoc pairwise comparisons were conducted by averaging across the levels of age and ROIs. Paired two-tailed t-tests were performed within each ROI to compare differences between hemispheres within each cross-condition direction. Cross-condition decoding was higher in the left hemisphere compared to the right hemisphere for both cross-condition direction: shallow-to-deep [Left: Mean =.531 SE = 3.8e-3, Right: Mean = .513 SE = 3.3e-3, t(54) = 5.553, p < .001] and deep-to-shallow [Left: Mean = .521 SE = 2.6e-3, Right: Mean = .510 SE = 2.3e-3, t(54) = 5.060, p < .001] (Figure S3).

**Figure S2.**
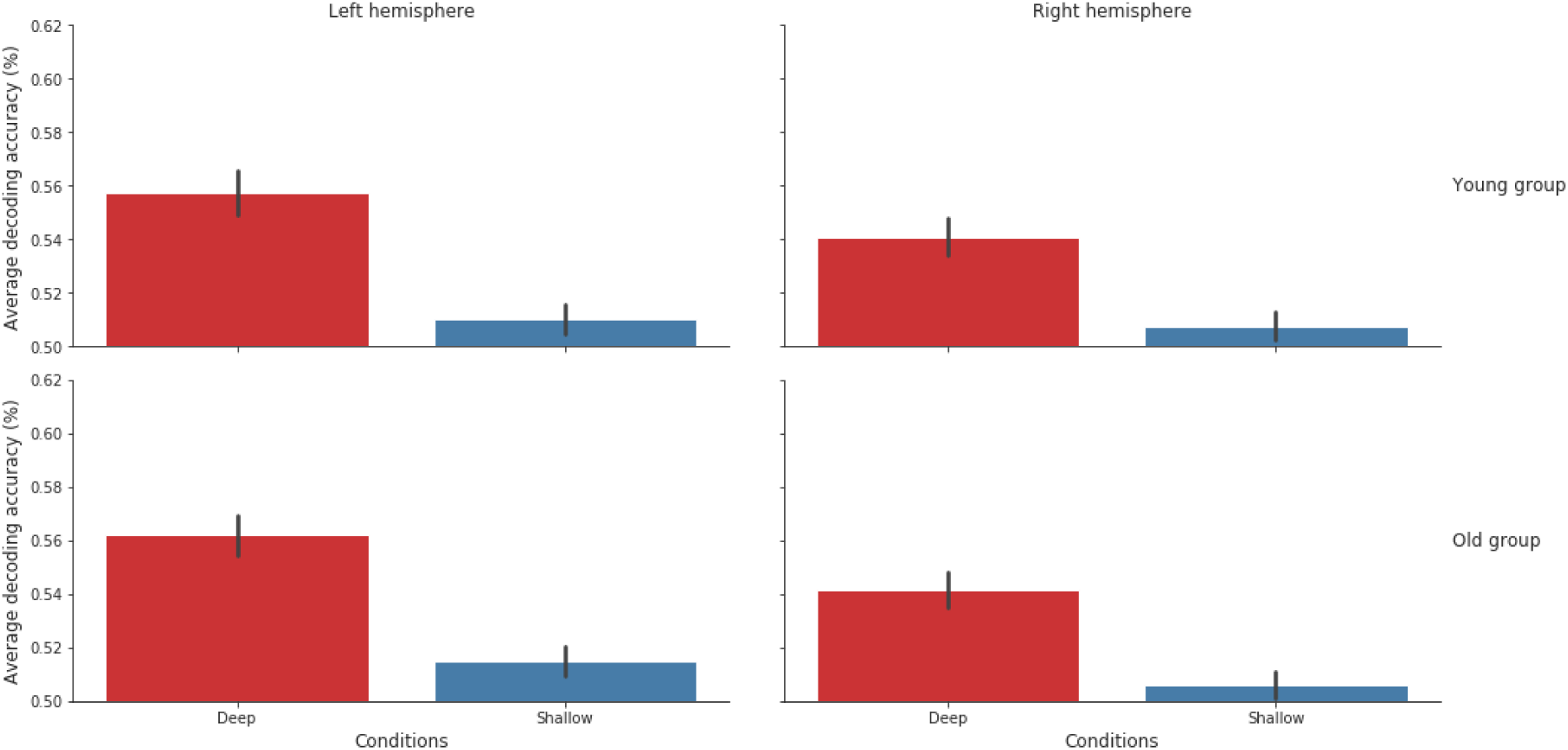
Average decoding accuracy obtained from the within condition analysis by hemisphere, age-group, and condition. The error bars represent a 95% confidence interval.

**Figure S3.**
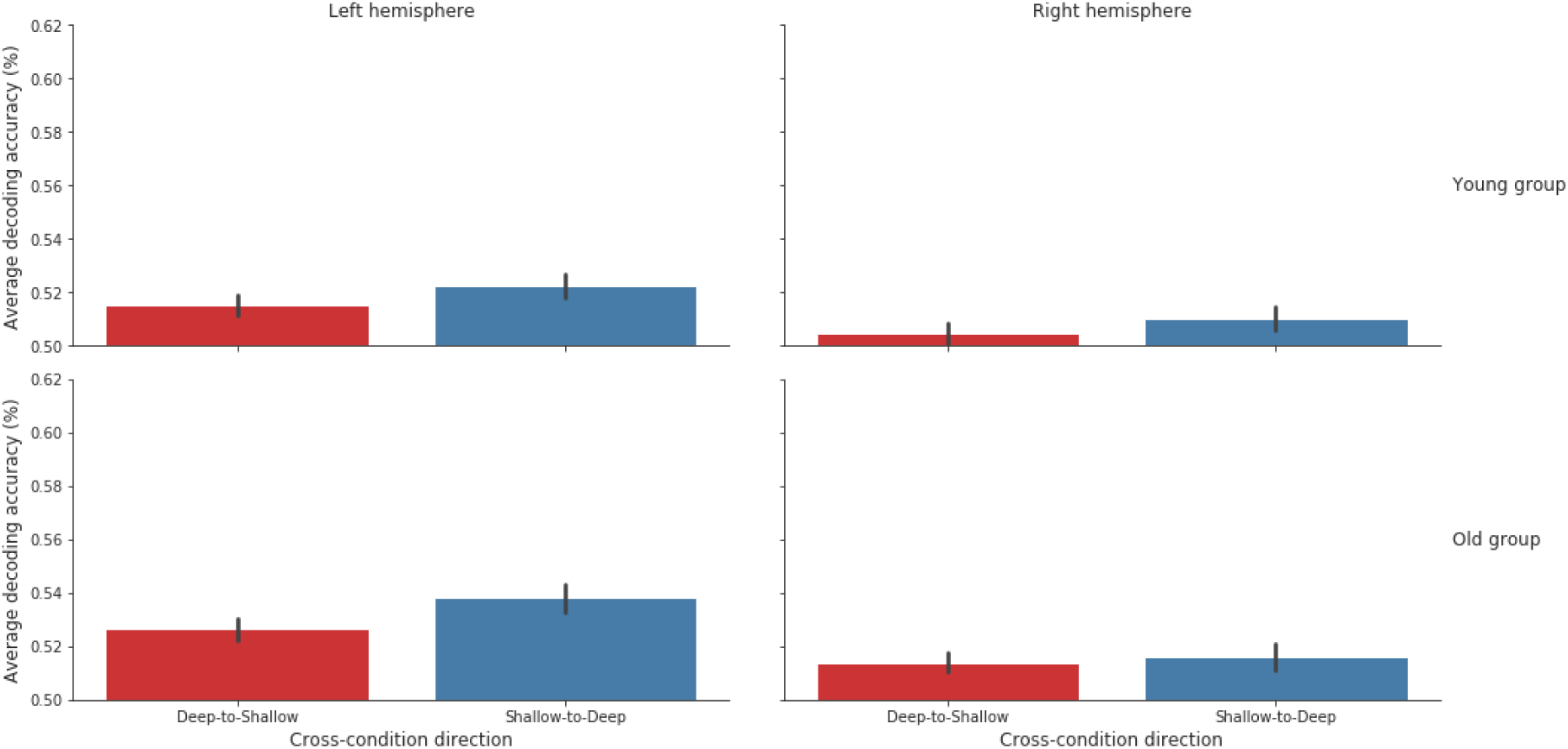
Average decoding accuracy obtained from the cross-condition analysis by hemisphere, age-group, and condition. The error bars represent a 95% confidence interval.

#### Greater decoding accuracies are observed in the deep compared to the shallow condition in the context of within condition decoding analyses

In the within-condition decoding analysis, we observed a significant main effect of condition [F(1, 54) = 18.553, p < .001, 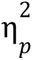 = .256]. Decoding accuracies were significantly higher in the deep (Mean = .551, SE = .007) compared to the shallow condition (Mean = .510, SE = .007) (Figure S2). Additionally, we found interactions between the condition factor and other variables: condition * ROIs [F(6.328, 341.730) = 8.794, p < .001, 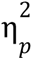 = .14], condition * hemisphere [F(1, 54) = 15.975, p < .001, 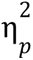 = .228], and condition * hemisphere * ROIs [F(11.927, 644.039) = 2.342, p = .011, 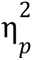 = .042]. As mentioned in the hemisphere and interaction effects section, we aimed to further explore the hemisphere * condition * ROIs interaction observed in the within-condition decoding analysis. In the following paragraph, our focus is on examining how decoding accuracies differ between conditions within each hemisphere for each ROI.

To investigate this, we conducted two additional mixed-effects ANOVA analyses, one for the left hemisphere and another for the right hemisphere, using 15 ROIs, 2 conditions, and 2 age groups as factors. The DOF for these ANOVAs were corrected using the Huynh-Feldt method. In both the left [F(1, 54) = 24.346, p < .001, 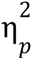 = .311] and right [F(1, 54) = 12.736, p < .001, 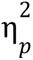 = .191] hemisphere ANOVAs, we found a significant main effect of condition. Overall, decoding accuracies were higher in the deep condition [Left: Mean = .560, SE = .007; Right: Mean = .541, SE = .007] compared to the shallow condition [Left: Mean = .513, SE = .007; Right: Mean = .507, SE = .007] (Figure S2). Furthermore, we observed an interaction between condition and ROIs in both the left [F(7.436, 401.568) = 9.261, p < .001, 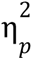 = .146] and right [F(8.830, 476.797) = 4.062, p < .001, 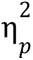 = .070] hemispheres. To explore these interactions, pairwise comparisons were performed within each hemisphere, comparing the shallow and deep conditions in each ROI while averaging over the levels of age. Post-hoc paired two-tailed t-tests with p-values corrected using the FDR for multiple comparisons were conducted.

In the left hemisphere, decoding accuracies were significantly higher in the deep condition compared to the shallow condition in almost all ROIs, except for the middle orbitofrontal gyrus [Deep: Mean = .530 SE = 8.5e-3, Shallow: Mean = .516, SE = 7.9e-3, t(54) = 1.203, p = .233] and the temporal pole [Deep: Mean = .518 SE = 7.3e-3, Shallow: Mean = .502, SE = 7.3e-3, t(54) = 1.348, p = .196]. Similarly, in the right hemisphere, decoding accuracies were significantly higher in the deep condition compared to the shallow condition in all ROIs except for the parahippocampal gyrus [Deep: Mean = .522 SE = 9.7e-3, Shallow: Mean = .507 SE = 6.8e-3, t(54) = 1.210, p = .248], posterior cingulate [Deep: Mean = .536 SE = 9.4e-3, Shallow: Mean = .514 SE = 7.4e-3, t(54) = 1.980, p = .061], and temporal pole [Deep: Mean = .504 SE = 6.6e-3, Shallow: Mean = .501 SE = 7.3e-3, t(54) = .274, p = .785] (Figure 3).

#### Asymmetries in the cross-condition direction generalization

We also note that the analysis of cross-condition decoding accuracies using a 15 (ROIs) * 2 cross-condition direction * 2 hemisphere * 2 age mixed-effects ANOVA with Huynh-Feldt corrected DOF revealed significant asymmetries in decoding directionality (van den Hurk & Op de Beeck, 2019). The results showed that decoding accuracies were higher when the decoder was trained on the shallow processing condition and tested on the deep processing condition (i.e., shallow-to-deep; Mean = .522, SE = .003), compared to the reverse direction (i.e., deep-to-shallow; Mean = .515, SE = .003) [F(1, 54) = 9.557, p = .003, 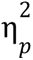 = .150] particularly in the left hemisphere (Figure S3). This finding appears counterintuitive given that we did not observe significant decoding of semantic categories above chance level within the shallow condition, and it remains unclear how the trained classifier in the shallow condition could generalize effectively to the deep condition. Furthermore, considering the high decoding accuracy in the deep condition, one might have expected better generalization from deep-to-shallow rather than shallow-to-deep. Similar asymmetries in decoding directionality have been reported in previous studies (e.g., Akama et al., 2012; Johnson & Johnson, 2014; Quadflieg et al., 2011; van den Hurk et al., 2017). Building on the work of van den Hurk & Op de Beeck (2019), we propose that these cross-classification decoding asymmetries may not be directly linked to experimental or neurobiological factors, but rather to differences in SNR ratios between the datasets of shallow processing and deep processing conditions. Van den Hurk & de Beeck demonstrated that a dataset with higher SNR yielded good within-condition classifications but poor generalization to a dataset with lower SNR, and vice versa. They suggested that the orientation of a soft margin SVM hyperplane, trained on a dataset with high SNR, might be more sensitive to adaptations of support vectors in the test set, leading to poor generalization in datasets with higher noise levels. This trend has been observed in other studies as well (Akama et al., 2012; Johnson & Johnson, 2014; Quadflieg et al., 2011; van den Hurk et al., 2017), aligning with our own findings.

Despite the absence of interaction effects between cross-condition direction and age factors in our main ANOVA [F(1, 54) = .015, p = .904], we conducted an additional ANOVA with 15 ROIs * 2 hemispheres * 2 age groups as factors specifically in the deep-to-shallow cross-classification direction in order to address potential biases in our interpretations due to SNR. Importantly, we observed a main effect of age (F(1, 54) = 5.613, p = .021, 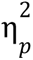 = .094), indicating that deep-to-shallow decoding was higher in the older group (Mean = .520, SE = .003) compared to the young group (Mean = .510, SE = .003) (Figure S4).

**Figure S4.**
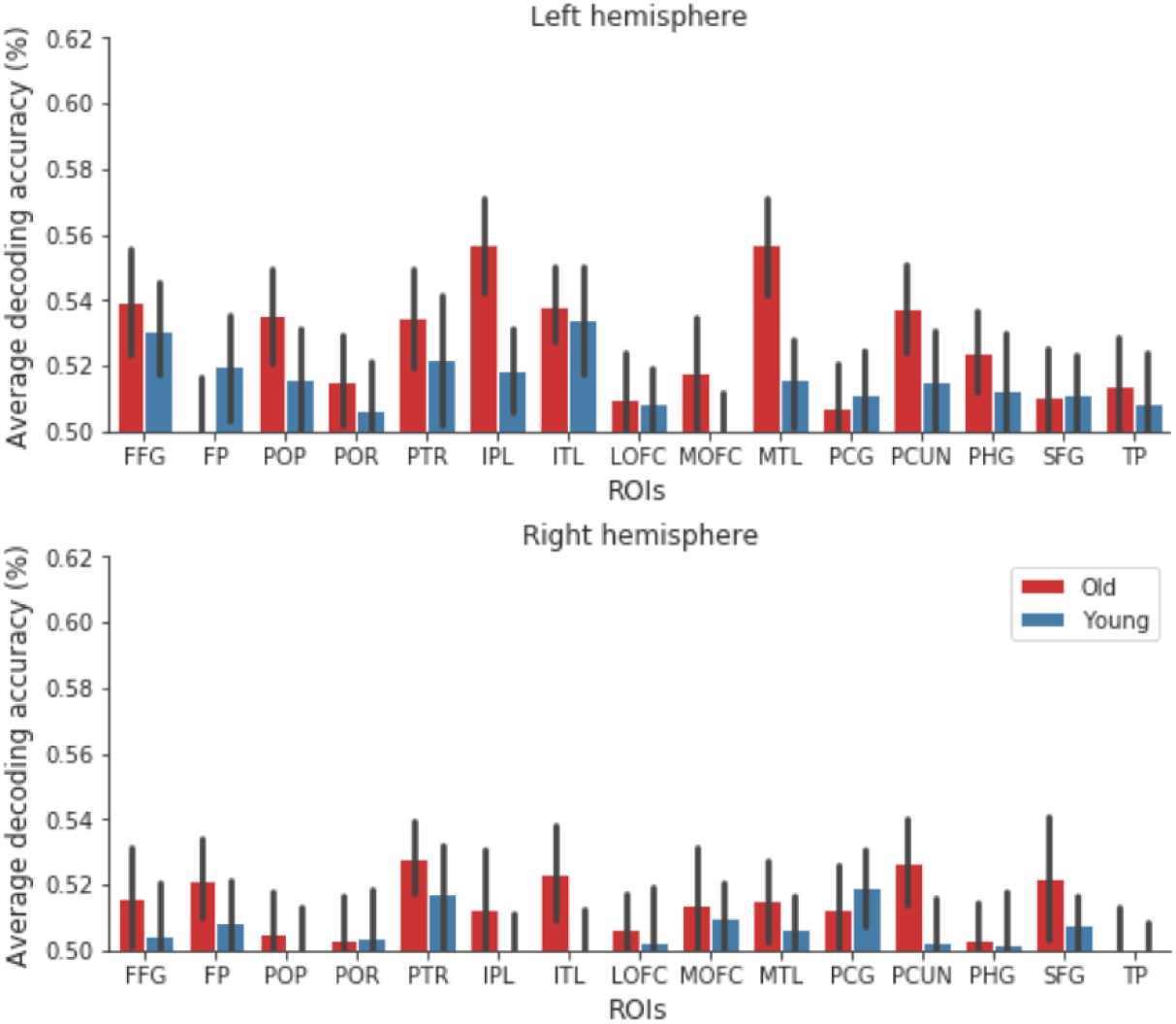
Cross-condition decoding accuracies in deep-to-shallow condition by ROI, hemisphere, and age-group. The error bars represent a 95% confidence interval.

#### The inferior parietal lobe, middle temporal gyrus and inferior temporal gyrus exhibited the highest decoding accuracies in both within-condition and cross-condition analyses

Given that the previous sections have already provided detailed information on the interactions between ROIs and other factors, we will now provide a brief overview of the overall decoding accuracy across the 15 ROIs, averaged over all levels of other factors (i.e., condition or cross-condition, hemisphere, age). In the within-condition decoding analysis, we observed a significant main effect of ROIs [F(4.682, 253.377) = 2.145, p < .001, 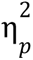 = .205]. The ROIs with the highest decoding accuracies were the inferior parietal lobe (Mean = .552, SE = 7e-3), inferior temporal gyrus (Mean = .545, SE = 6.7e-3), middle temporal gyrus (Mean = .543, SE = 6.5e-3), and fusiform gyrus (Mean = .540, SE = 6.8e-3). On the other hand, the ROIs with the lowest decoding accuracies were the parahippocampal gyrus (Mean = .517, SE = 5.1e-3), temporal pole (Mean = .506, SE = 3.7e-3), and frontal pole (Mean = .506, SE = 4.4e-3) (Figure 3).

In the cross-condition decoding analysis, we observed a significant main effect of ROIs [F(4.682, 253.377) = 2.145, p < .001, 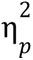 = .205]. The ROIs with the highest cross-condition decoding accuracies were the fusiform gyrus (Mean = .540, SE = 5e-3), inferior parietal lobe (Mean = .530, SE = 5.2e-3), middle temporal gyrus (Mean = .529, SE = 4.6e-3), and inferior temporal gyrus (Mean = .528, SE = 4.3e-3). Conversely, the ROIs with the lowest cross-condition decoding accuracies were the frontal pole (Mean = .509, SE = 2.8e-3), lateral orbitofrontal gyrus (Mean = .507, SE = 3.9e-3), and temporal pole (Mean = .506, SE = 3.3e-3) (Figure 4).

